# The conformational landscape of a serpin N-terminal subdomain facilitates folding and in-cell quality control

**DOI:** 10.1101/2023.04.24.537978

**Authors:** Upneet Kaur, Kyle C. Kihn, Haiping Ke, Weiwei Kuo, Lila M. Gierasch, Daniel N. Hebert, Patrick L. Wintrode, Daniel Deredge, Anne Gershenson

## Abstract

Many multi-domain proteins including the serpin family of serine protease inhibitors contain non-sequential domains composed of regions that are far apart in sequence. Because proteins are translated vectorially from N-to C-terminus, such domains pose a particular challenge: how to balance the conformational lability necessary to form productive interactions between early and late translated regions while avoiding aggregation. This balance is mediated by the protein sequence properties and the interactions of the folding protein with the cellular quality control machinery. For serpins, particularly α_1_-antitrypsin (AAT), mutations often lead to polymer accumulation in cells and consequent disease suggesting that the lability/aggregation balance is especially precarious. Therefore, we investigated the properties of progressively longer AAT N-terminal fragments in solution and in cells. The N-terminal subdomain, residues 1-190 (AAT190), is monomeric in solution and efficiently degraded in cells. More β-rich fragments, 1-290 and 1-323, form small oligomers in solution, but are still efficiently degraded, and even the polymerization promoting Siiyama (S53F) mutation did not significantly affect fragment degradation. *In vitro,* the AAT190 region is among the last regions incorporated into the final structure. Hydrogen-deuterium exchange mass spectrometry and enhanced sampling molecular dynamics simulations show that AAT190 has a broad, dynamic conformational ensemble that helps protect one particularly aggregation prone β-strand from solvent. These AAT190 dynamics result in transient exposure of sequences that are buried in folded, full-length AAT, which may provide important recognition sites for the cellular quality control machinery and facilitate degradation and, under favorable conditions, reduce the likelihood of polymerization.

## Introduction

Globular protein folding is key to function, and while recent advances in protein structure prediction have revealed the fold of hundreds of thousands of proteins, the process by which proteins fold is still not well understood. Protein folding studies have focused primarily on small (100-150 amino acid) single domain proteins (1, 2); however, many proteins have multiple domains which can complicate folding. Multi-domain proteins may contain sequential domains, “beads-on-a-string”, where domains are translated one after another from the N-to the C-terminus, non-sequential domains, where domains contain regions that are far apart in the sequence, or some combination of the two. While all proteins must avoid aggregation during folding (3), this avoidance may be particularly troublesome for proteins with non-sequential domains because as the protein is translated N-terminal regions of a non-sequential domain must both avoid oligomerization/aggregation and maintain enough flexibility to properly assemble with pieces that are translated later. This balance between the need to populate a larger, more diverse conformational ensemble during folding while avoiding oligomerization is mediated both by the protein sequence and by the cellular protein quality control machinery.

One way to investigate this balance is to mimic the vectorial nature of protein translation by the ribosome by expressing progressively longer N-terminal fragments of a protein both as purified fragments and in cells. The purified fragments provide data on the intrinsic conformational and oligomerization propensities of the sequence (e.g., (4–8)) while in-cell studies can help reveal how the cellular environment and protein quality control machinery modulate the probability of oligomerization and whether aberrant oligomers are properly targeted for degradation. Members of the serpin structural superfamily are ideal for studying how non-sequential multi-domain proteins balance flexibility and oligomerization propensity. Serpins consist of two non-sequential domains (9) (Fig. 1), and many mutations in secretory serpins lead to pathological oligomerization in the endoplasmic reticulum (ER) (e.g., (10–16)). Thus, the comparison between purified fragments and in-cell behavior is particularly biologically relevant.

**Figure 1.**
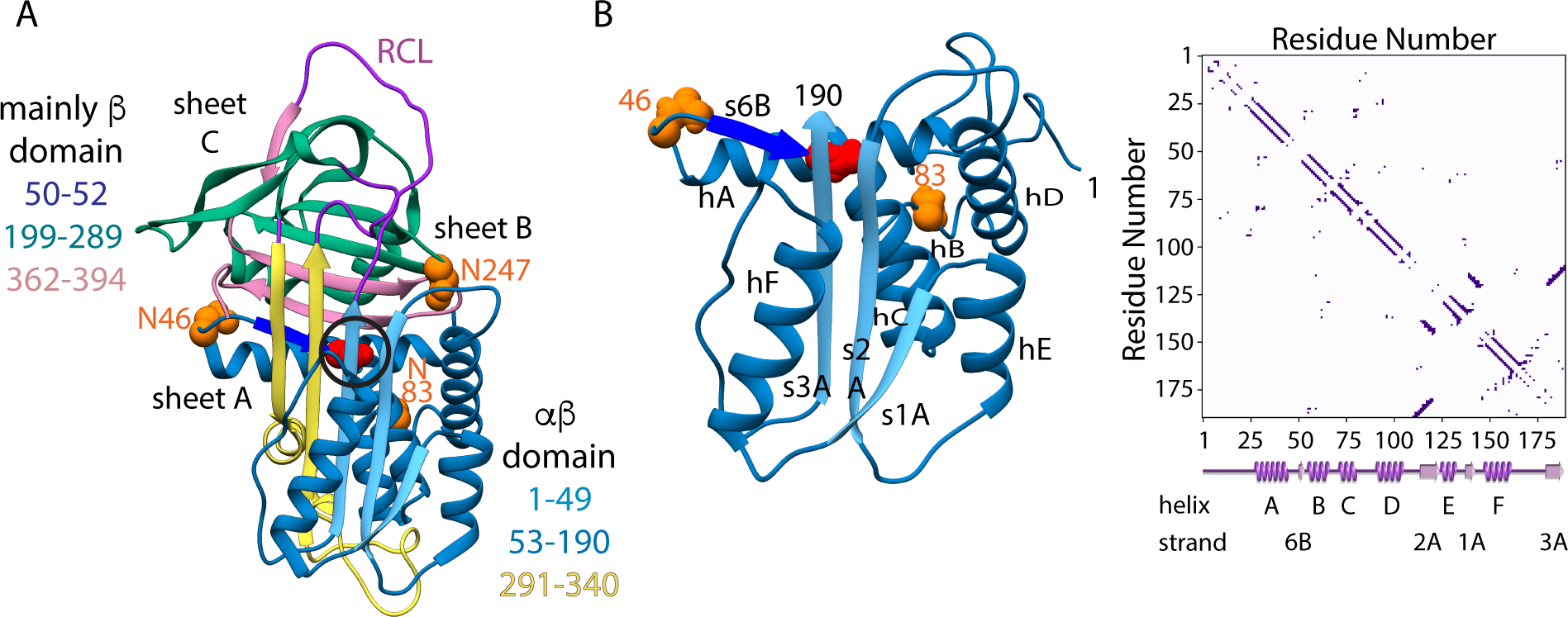
AAT structure showing the complex serpin topology. (A) The x-ray crystal structure of full-length AAT (PDB:1QLP (36)). The N-and C-terminal regions of the αβ domain are in blue and yellow, respectively; the N-terminal regions of the mainly β domain are in bright blue (strand 6B) and green with the C-terminal region in pink; loops connecting the two domains, including the long functionally required RCL, are in purple; and asparagines that are glycosylated in the ER are in orange spacefill. The black circle indicates the functionally relevant shutter region containing the Siiyama (S53F) mutation (red spacefill). (B) Structure and contact map for the 1 to 190 fragment (AAT190) in the context of the full-length active protein. The contact map includes the disordered N-terminus (residues 1-22) which was added using the CHARMM-GUI (37). The contact map for the entire full-length protein and structures of the longer fragments, 1-290 (AAT290) and 1-323 (AAT323), in the context of the full-length protein are shown in Figure S1.

Serpins adopt a topologically complex, ellipsoidal fold consisting of two non-sequential domains (9), an αβ two layer sandwich characterized by the central β-sheet A that defines the long axis of the ellipsoid plus a smaller mainly β domain (Fig. 1). Perhaps the best-studied member of the superfamily is the 394 amino acid long secretory and inhibitory serpin α_1_-antitrypsin (AAT). AAT is abundant in human plasma where it inhibits proteases associated with inflammation, and its equilibrium and kinetic folding have been extensively studied (17–27). As a secretory protein AAT folds and matures in the ER, and AAT mutations that lead to accumulation of AAT polymers in the ER of liver cells (the main source of circulating AAT) cause both gain-of-function liver disease due to polymer accumulation and loss of function lung disease due to low circulating AAT levels (28–31).

We therefore studied the behavior of progressively longer AAT N-terminal fragments *in vitro* and in Chinese hamster ovary (CHO) cells. The shortest fragment, encompassing residues 1-190 (AAT190), was monomeric in solution, in agreement with recent studies by Cabrita and co-workers (32), and was easily degraded by ER-associated protein degradation (ERAD) in CHO cells. By contrast, two longer fragments (AAT290 and AAT323) formed small oligomers in solution but were still readily disposed of by ERAD. Surprisingly, while full-length AAT containing the polymerization promoting mutation Siiyama (Ser53Phe) (33–35) accumulated in CHO cells as expected, AAT fragments containing this mutation were efficiently recognized and targeted for degradation. Thus, even for polymerization promoting mutations located early in the sequence, the full-length protein with an intact C-terminus appears to be key to pathological polymerization.

The finding that AAT190 is soluble and monomeric in solution agrees with results from both experiments (32) and simulations of AAT folding (26, 27, 38), which suggest that structural consolidation of the N-terminal portion of the αβ domain is one of the last steps in folding of full-length AAT. This accumulation of evidence led us to hypothesize that AAT190 could serve as a convenient handle for recognition by protein quality control. In order to gain insights into the conformational ensemble that might provide such a handle, we generated likely AAT190 conformational ensembles by combining hydrogen-deuterium exchange mass spectrometry (HDX-MS) of purified AAT190 with enhanced sampling molecular dynamics (MD) simulations (39, 40). The resulting diverse conformational ensembles showcase how AAT190 shields a particularly oligomerization prone β-strand while also being conformationally labile, a property that likely aids in recognition by ER quality control.

## Results

### AAT structure and conformational lability

The non-sequential serpin αβ and mainly β domains are connected by three linkers (Fig. 1), the last of which, the solvent-exposed reactive center loop (RCL) near the C-terminus is key to function for inhibitory serpins (15, 41, 42). Docking of a target serine or cysteine protease on the RCL leads to the formation of an acyl-enzyme bond between the catalytic serine (cysteine) of the protease and the scissile bond of the RCL. Like the springing of a mousetrap, subsequent RCL cleavage initiates insertion of the N-terminal portion of the cleaved RCL as β-strand 4 in the central β-sheet A translocating the covalently linked protease approximately 70 Å across the serpin, mechanically disrupting the protease and resulting in a kinetically trapped protease-serpin complex (15, 17, 41, 42). The energy required for mechanical inhibition of the protease is stored in the metastable active serpin conformation. A number of AAT mutations lead to accumulation of AAT polymers in the ER (11, 12) and this polymerization propensity may be related to the requirement that the sequence of AAT and other inhibitory serpins encode the ability to expand the central β-sheet.

### AAT N-terminal fragments from monomers to oligomers

AAT N-terminal fragments that are likely to fold autonomously were identified based on the results of AAT folding simulations (26, 27), subdomain boundaries as defined as by the locations of loops connecting the two serpin domains, and the prediction of likely autonomous folding units using the Rapid Autonomous Fragment Test (RAFT) methodology (43). All of these analyses and subsequent experiments used the M1V human AAT sequence, the most common wild-type sequence (44). The RAFT analysis suggests the N-terminal subdomain of the αβ domain (residues 23-190) has the highest probability of forming autonomous local structure (Fig. S2). High RAFT scores indicate that there are more contacts between residues within a given fragment than there are between residues in the fragment and residues in the rest of the protein (for details see the Experimental Procedures), but RAFT scores do not report on the stability of contacts. The high RAFT score for the N-terminal subdomain of the αβ domain is therefore consistent with all-atom folding simulations (26) suggesting that local structure forms in this same region and a recent report that the isolated 1 to 191 fragment from AAT forms a molten-globule like structure (32). The fragment containing residues 91 to 289 also has high RAFT scores consistent with the experimentally observed early folding of residues 200-290, the N-terminal subdomain of the mainly β domain (24). These considerations led us to investigate the structure of the N-terminal αβ subdomain AAT190 (residues 1-190) and AAT290 (residues 1-290) which encompasses the N-terminal subdomains of both the αβ and mainly β domains. As a secretory protein, AAT folds and matures in the ER, and we therefore also investigated the structure of AAT323 (residues 1-323), which, assuming that the ribosome tunnel and ER translocon sequester 70 amino acids (45, 46), is one of the longest AAT ribosome-attached nascent chains in the ER lumen.

To avoid disulfide bond formation in solution, the single cysteine residue (C232) in AAT was mutated to serine (see Table S1 for fragment amino acid sequences). The fragments were expressed in *Escherichia coli*, isolated as inclusion bodies and characterized following refolding and removal of the N-terminal 6XHis-tag. AAT190 showed significant amounts of secondary structure by far UV circular dichroism (CD) (Fig. 2A), and sedimentation velocity analytical ultracentrifugation (AUC) experiments revealed that AAT190 is monomeric (Fig. 2B). These results are in agreement with recent CD and NMR results from Plessa *et al.* showing that the AAT N-terminal fragment containing residues 1-191 has secondary structure and is monomeric (32). In contrast, the two longer fragments AAT290 (Fig. 2C) and AAT323 (Fig. 2D) formed small oligomers with the number of monomers per oligomer increasing with fragment length. Thus, the oligomerization propensity of the AAT fragments increases as a function of chain length perhaps due to a corresponding increase in the number of unsatisfied β-strands.

**Figure 2.**
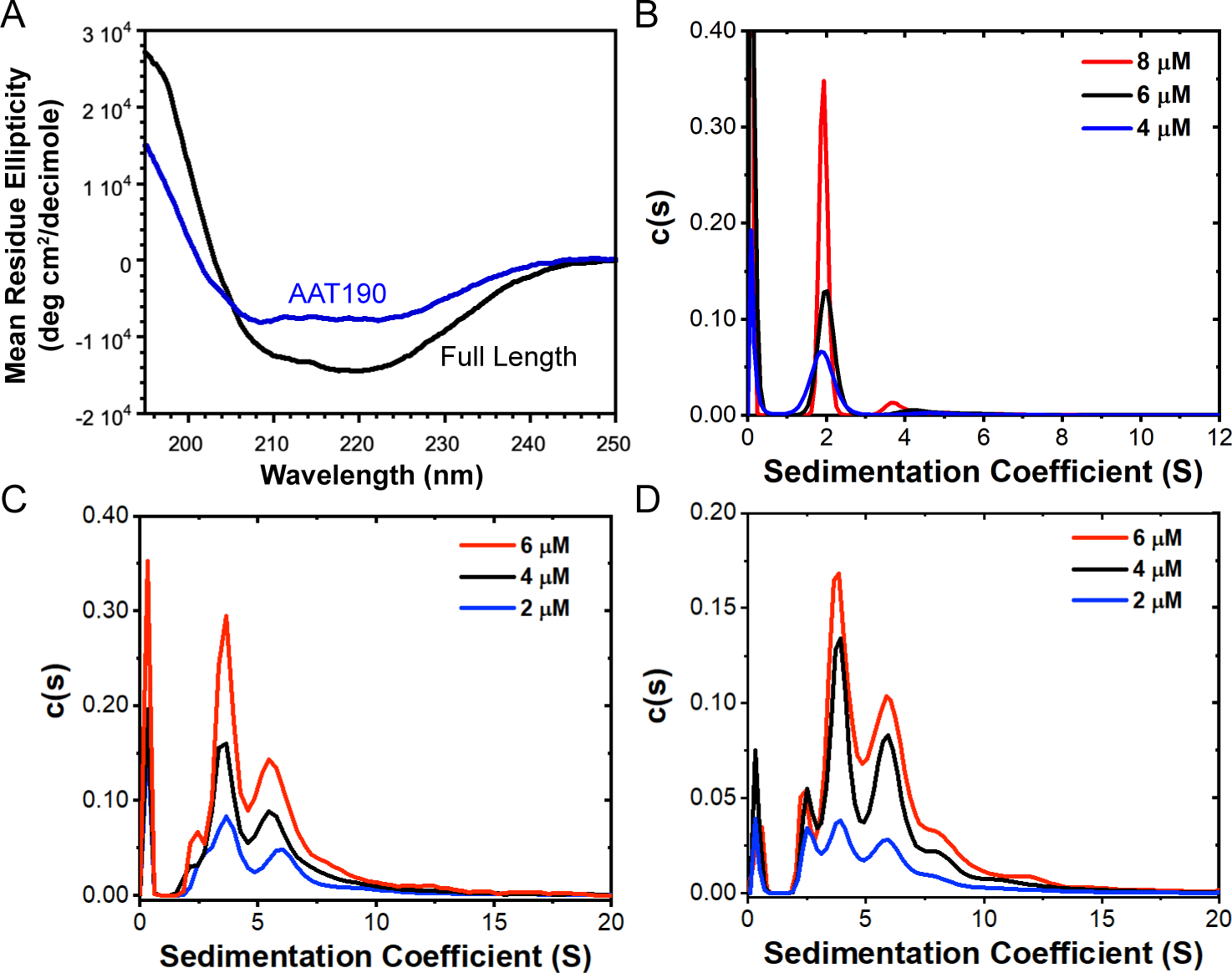
AAT190 is monomeric while longer fragments form small oligomers. (A) Far UV circular dichroism (CD) spectra for AAT190 (blue) and full-length AAT (black) show that AAT190 has significant amounts of secondary structure. (B) Distributions of sedimentation coefficients for 8 (red), 6 (black) and 4 μM of AAT190 indicating that this fragment is mainly monomeric. Distributions of sedimentation coefficients for 6 (red), 4 (black) and 2 μM (blue) of (C) AAT290 and (D) AAT323 showing mainly dimers for both AAT290 and AAT323, and that more higher order oligomers are populated for AAT323. For all distributions, the peak near 0 arises from a population of small molecules much smaller than the protein fragments. For AUC data, fits and residuals see Figure S3 for AAT190 and Figure S4 for AAT290 and AAT323.

### The cellular fate of AAT fragments

A number of full-length disease-associated AAT mutants accumulate as polymers in the ER and are associated with cell death (10, 14, 47). One antibody that recognizes mutant polymers also recognizes polymers formed by heating purified wild-type AAT suggesting that polymer formation is at least partially encoded in the wild-type sequence (48). Our observations that the oligomerization propensity of AAT N-terminal fragments increases as a function of chain-length led us to ask whether the cell’s ability to clear AAT oligomers might also vary with chain length. We therefore expressed the AAT fragments and full-length AAT in CHO cells and characterized their fate (Fig. 3).

**Figure 3.**
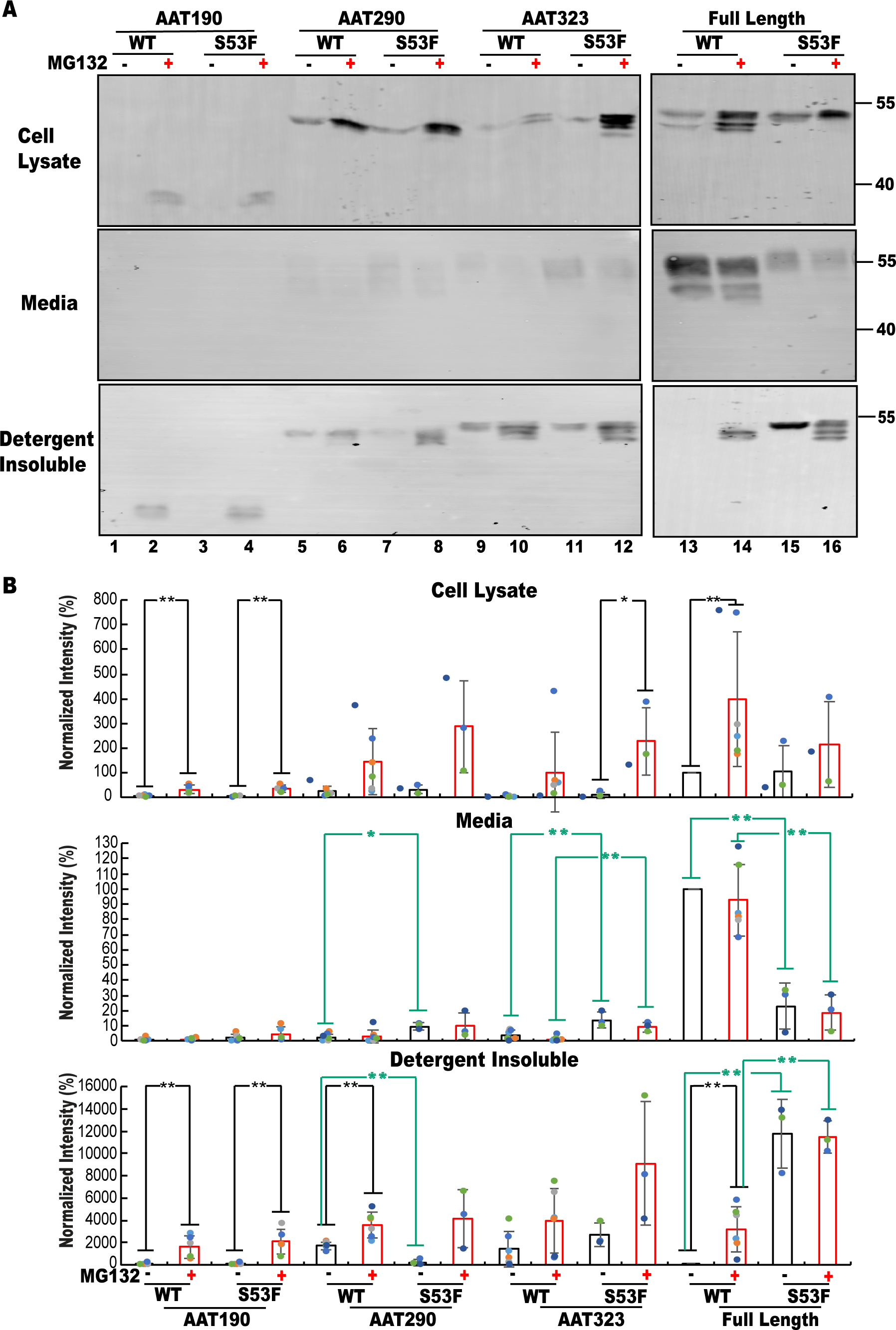
AAT N-terminal fragments are targeted for degradation, and fragment trafficking is only slightly perturbed by the Siiyama (S53F) mutation. (A) CHO cells were transfected with AAT N-terminal fragments or full-length M1V AAT without (wild-type, WT) or with the Siiyama mutation (S53F). The cells were grown for 20 hr, then grown for a further 20 hr in the absence (black) or presence (red) of the proteasome inhibitor MG132. Proteins from the cell lysate, media and detergent (Triton X-100) insoluble fractions were precipitated with trichloroacetic acid (TCA), solubilized and resolved by reducing 9% SDS-PAGE. AAT constructs were detected using an anti-Myc antibody. (B) Quantification of the results in A from three to five biological replicates are shown as colored dots. The absence or presence of MG132 is indicated by black and red bars, respectively. For each fraction and replicate the fluorescence intensities from the immunoblots were normalized to the intensity of full-length wild-type AAT (M1V) in the absence of MG132. A single asterisk indicates p-values<0.05 while a double asterisk indicates p-values<0.01 from unpaired t-tests. Comparisons between +/-the proteasome inhibitor MG132 and between the M1V and Siiyama backgrounds are indicated by black and green lines, respectively.

To ensure ER targeting and to monitor cellular expression, the AAT signal sequence and a Myc tag were added to each fragment construct (Table S2). The steady-state fraction of the fragment or full-length AAT population that was successfully secreted from the cell (media), retained in the cell as a soluble protein (cell lysates) or aggregated (detergent-insoluble) was determined by immunoblotting using an anti-Myc antibody (Fig. 3). Active N-terminally tagged full-length wild-type AAT was efficiently expressed and secreted from CHO cells as expected (Fig. 3A, lane 13 and Fig. 3B). All of the fragments were largely retained inside the cells. AAT190 was observed at low levels with little accumulation in the detergent-insoluble fraction consistent with its presumably monomeric character (Fig. 3A lane 1 and Fig 3B). AAT290 and AAT323, were detected at higher levels in the cell lysate and, as expected from the oligomerization propensity of the purified fragments, significant amounts of these constructs accumulated in the detergent-insoluble fraction (Fig. 3A lanes 5 and 9, and Fig. 3B). Their cellular retention indicates that the ER quality control network appropriately recognized these fragments as incompletely or improperly folded, consistent with the finding that AAT N-terminal fragments with lengths ranging from 334 to 391 amino acids were not secreted from Cos-1 cells (49).

Prolonged ER retention can result in protein degradation (50). While retained proteins are often cleared by the proteasome through the ERAD pathway (50), polymers formed by full-length AAT mutants may be ERAD resistant resulting in the activation of other pathways (51, 52). To determine whether the AAT fragments were being degraded by ERAD, cells were treated with the proteasome inhibitor MG132. MG132 treatment significantly increased the detected cellular levels for all AAT constructs including the full-length protein (Fig. 3A lanes 13 and 14).

AAT190 is barely detected under normal experimental conditions and proteasome inhibition leads to a particularly significant increase in detectable product (Fig. 3A lanes 1 and 2, and Fig. 3B), indicating that the shortest fragment was efficiently cleared by ERAD. Further support for the importance of ERAD in the degradation of AAT fragments was provided by comparisons between the fate of AAT323 and the classic ERAD substrate AAT *null* Hong Kong (NHK), which has a frame shift at codon 318 resulting in a 333 residue-long truncated AAT variant composed of residues 1 to 318 followed by a 15 amino acid long missense peptide and a stop codon ((53),Table S2). Radioactive pulse-chase analysis demonstrated that the time course of ER retention, turnover and aggregation of AAT323 was similar to NHK (Fig. S5). AAT323 is 98% identical to NHK, and the similarities in their fates and importance of ERAD for cellular clearance of both fragments suggest that the AAT sequence rather than the missense tail dictates the cellular fate of NHK.

For all the AAT constructs, MG132-mediated proteasome inhibition also significantly increased the amount of protein in the detergent-insoluble fraction, demonstrating that the ER-retained protein was highly susceptible to aggregation. While this oligomerization propensity does not significantly impair cellular clearance of wild-type fragments and full-length protein, it does resemble the pathogenic polymerization and accumulation of disease-associated mutants such as Z (Glu342Lys), Mmalton (τιF52) and Siiyama (S53F) in the ER (11, 33–35). While Z is a common and well-characterized pathogenic mutant, much less is known about mutations near the N-terminus that are associated with AAT polymerization. We therefore investigated the fate of fragments containing the Siiyama mutation. Siiyama-containing AAT190 (AAT190S) was expressed in both *E. coli* and CHO cells. Attempts to solubilize *E. coli* AAT190S inclusion bodies were unsuccessful. This inability to solubilize inclusion bodies is similar to what is observed when expressing full-length oligomerization-prone AAT mutants such as Z in *E. coli* (54), and we therefore chose not to further pursue *E. coli* expression and purification of Siiyama-containing AAT variants.

As expected (55, 56), expression of full-length AAT containing the Siiyama mutation in CHO cells led to decreased secretion and intracellular accumulation when compared to the wild-type full-length protein (Fig. 3A, lanes 13 and 15 and Fig. 3B). A significant amount of the full-length Siiyama AAT variant accumulated in the detergent-insoluble fraction. For the fragments, the poor expression of AAT190S was similar to that of the corresponding wild-type fragment (Fig. 3A, lanes 1and 3). Unexpectedly however, the longer fragments containing the Siiyama mutation (AAT290S and AAT323S) showed significant increases in secretion (Fig. 3A, lanes 5, 7, 9 and 11), and less of the AAT290S fragment accumulated in the detergent-insoluble fraction when compared to AAT290 (Fig. 3B). Protease inhibition by MG132 increased the amount of Siiyama fragments in both the cell lysate soluble and detergent-insoluble fractions, indicating that the Siiyama fragments were efficiently degraded by ERAD. In contrast, full-length AAT containing the Siiyama mutation was not efficiently targeted for ERAD and accumulated in cells and in detergent-insoluble fractions in the presence or absence of MG132. Overall, as observed for the purified wild-type fragments, the longer cellular expressed fragments were more oligomerization prone than the shorter AAT190 and this difference was augmented by the Siiyama mutation. Nonetheless, while AAT190S behaves differently in *E. coli*, the majority of fragments containing the Siiyama mutation are properly targeted for degradation in CHO cells.

### AAT190 Conformational Plasticity

AAT190 distinguishes itself from the other fragments by its monomeric character and the apparent ease with which it is targeted for ERAD. Based on both the far-UV CD spectrum (Fig. 2) and recently published investigations of the 1-191 AAT fragment (32), AAT190 likely has a molten globule character. While this characterization is informative, further details of the specific structural features present in the AAT190 conformational ensemble may provide clues to its recognition by the ER quality control network. We therefore used the HDXer methodology (39, 40), which combines HDX-MS and enhanced sampling MD simulations, to characterize the AAT190 conformational ensembles.

Pepsin digestion and liquid chromatography (LC)/MS, following HDX, identified a common set of 29 peptides covering 96.3% of AAT190 sequence (Fig. S6) enabling direct comparison of HDX data for AAT190 and full-length, active AAT. Most of the AAT190 structure was quite labile. Only 5 of the 29 peptides (14% percent of the AAT190 sequence) showed less than 50% exchange after 10 sec, and only 2 of the 5 protected peptides, 127-133 in helix E and 183-187 in strand 3A, showed 20% or lower exchange. Even this protection from exchange was mostly lost after only a 10 min incubation in deuterated solution (Fig. 4B). By contrast, in the context of full-length AAT (8 peptides) spanning 30% of the 1-190 fragment region, still displayed less than 50% exchange after 10 min of incubation (Fig. 4B). Nonetheless, the patterns of protection at 10 sec are similar in AAT190 and in the full-length protein (Fig. 4B and 4C).

**Figure 4.**
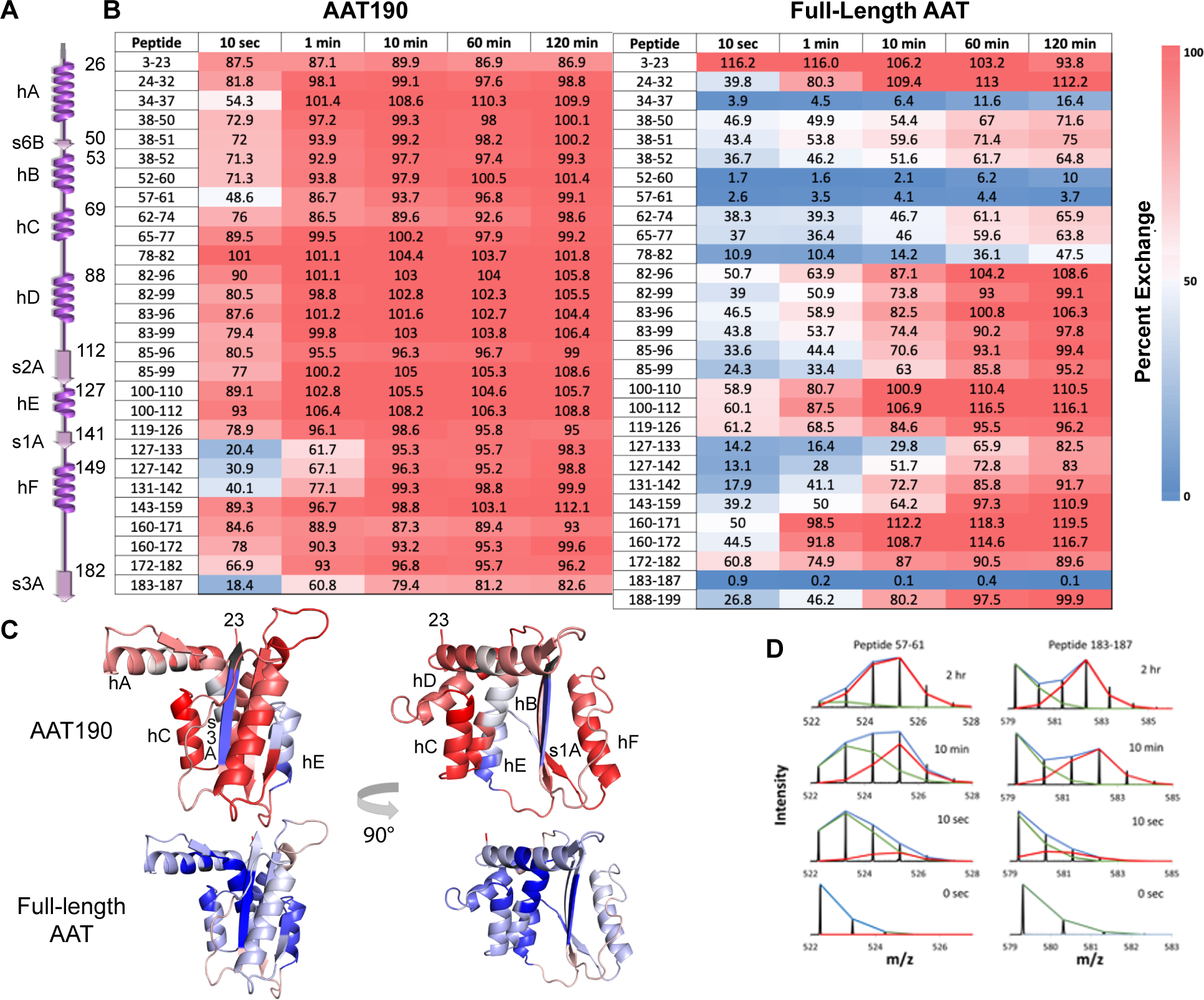
Structure in AAT190 is centered around strands 1 and 3A and helices A, B and E. (A) Schematic of the full-length AAT secondary structure. (B) Percent deuterium uptake for individual peptides at each incubation time point for AAT190 (left) and full-length AAT (right). Peptide sequence numbers are listed in the column on the left and percent deuteration over time is colored from blue (0%) to red (100%). Exchange time courses for each peptide are shown in Figure S7. (C) Percent exchange at 10 sec (0.167 min) for AAT190 (top) and residues 23-190 in the full-length protein (bottom). The percent exchange is mapped unto the structure of the fragment in the context of the full-length active protein structure (PDB: 1QLP (36)) colored as in B. Peptides not detected are shown in black. (D) Isotopic envelopes of peptides 57-61 (helix B) and 183-187 (strand 3A) showing the bimodal behavior characteristic of EX1 kinetics. As shown by AUC sedimentation velocity experiments, this AAT190 sample was mainly monomeric (Fig. S8). HDX-MS results for the AAT190 biological replicate and for all 394 amino acids in full-length AAT are shown in Figures S9 and S10, respectively.

In HDX-MS experiments two exchange mechanisms, EX1 and EX2, are generally observed. The less common EX1 exchange (57, 58) is characterized by bimodal isotopic mass envelopes. Such bimodal peaks can occur when a protein region undergoes slow cooperative unfolding/refolding with the lower mass peak corresponding to the protected species that has not yet undergone unfolding and the higher mass peak corresponding to species that have unfolded and undergone concerted HDX. In the more common EX2 mechanism (57), HDX is enabled by rapid small scale structural fluctuations resulting in a single isotopic envelope whose mass increases as a function of incubation time. In agreement with previous reports (59) all observed peptides in full-length AAT show only EX2 exchange. Surprisingly, in AAT190 EX1 exchange is observed for both the most protected peptide, 183-187 from strand 3A, where the protected envelope persisted well past 10 min and the less protected peptide 57-61 helix B peptide where the protected envelope persisted up to 1 min (Fig. 4D). In full-length AAT these same peptides are extremely well protected showing less than 5 percent exchange after 2 hr of incubation. These results suggest that: (i) most of the secondary and tertiary structure in AAT190, if present, is highly labile; (ii) regions in helices A, B and E along with strands 1A and 3A contain more stable structure; and (iii) surprisingly, the helix B and strand 3A regions may cooperatively unfold.

While the HDX-MS results provide important insights into AAT190 structural labilities, by themselves they do not provide full models of the conformational ensembles. Because the AAT190 conformational ensembles likely influence recognition by the ER quality control machinery, we used the HDX ensemble reweighting software package HDXer, recently developed by Bradshaw and coworkers (39, 40, 60) to elucidate likely AAT190 conformational ensembles. The HDXer approach was applied to AAT190 using the HDX-MS data and four simulated tempering MD simulations with different maximum temperatures which were combined into a single 200,000 frame trajectory. The combined HDXer (39, 60) and time-lagged independent component analysis (tICA) clustering (61, 62) resulted in statistically significant upweighting of six (out of 186 total) clusters which likely best reflect the HDX-MS results. As an independent test of the HDXer results, the frictional ratios, *f/fo*, were calculated for the six upweighted HDXer clusters using the HullRad webserver (63) and these results were compared to the *f/fo* values calculated from the AUC sedimentation velocity experiments. The frictional ratios calculated for the six upweighted HDXer clusters using the HullRad webserver (63) are generally higher than the average for the unweighted ensemble but are still less than *f/fo* calculated from the AUC sedimentation velocity experiments suggesting that the clusters may slightly overestimate AAT190 compaction (Fig. S11). This overestimation is expected because of known limitations in the force fields used for simulations of globular proteins (64–66). Nonetheless, because the upweighted conformational ensembles are consistent the HDX-MS results, they provide important insights into the location of transient secondary structure and the frequency of native versus non-native contacts.

While the resulting conformational ensembles are for purified AAT190, proteolysis of nascent full-length AAT chains (32) and AAT folding simulations (26) provide evidence that until late in folding, AAT190, i.e., the N-terminal region of the αβ domain, forms relatively few stable contacts with the rest of the protein. Therefore, the HDXer results for purified AAT190 provide plausible models for AAT190 conformational ensembles that might be populated in cells and recognized by ER quality control during AAT folding and maturation. The representative structures from each of the six clusters shown in Figures 5A and 5B highlight the conformational heterogeneity of AAT190. As can be seen from both the representative structures and the histograms of the radius of gyration (R_g_, Fig. 5C and 5D), the six clusters fall into two groups, with one group (clusters 1, 3 and 6) populating more expanded conformations while the other group (clusters 2, 4 and 5) is significantly more compact. This difference in compaction is also reflected in the relative persistence of native and non-native contacts as shown in the contact occupancy maps (Fig. 5E, 5F and S12). These maps display the frequency with which contacts are formed during the simulations, where points near the diagonal indicate contacts between residues close in sequence, while points further from the diagonal indicate contacts between sequentially distant residues. More persistent non-native contacts, likely indicating more persistent α-helical regions, lead to more compact structures and smaller R_g_s. While for simplicity the likely accessible conformations were grouped, in practice, AAT190 populates a diverse, dynamic conformational ensemble with interconversion between less compact and more compact conformations. Considering only the sterics of AAT190-chaperone interactions, it seems plausible to hypothesize that these less compact, more expanded AAT190 conformations may be more likely to be recognized by the ER quality control network.

**Figure 5.**
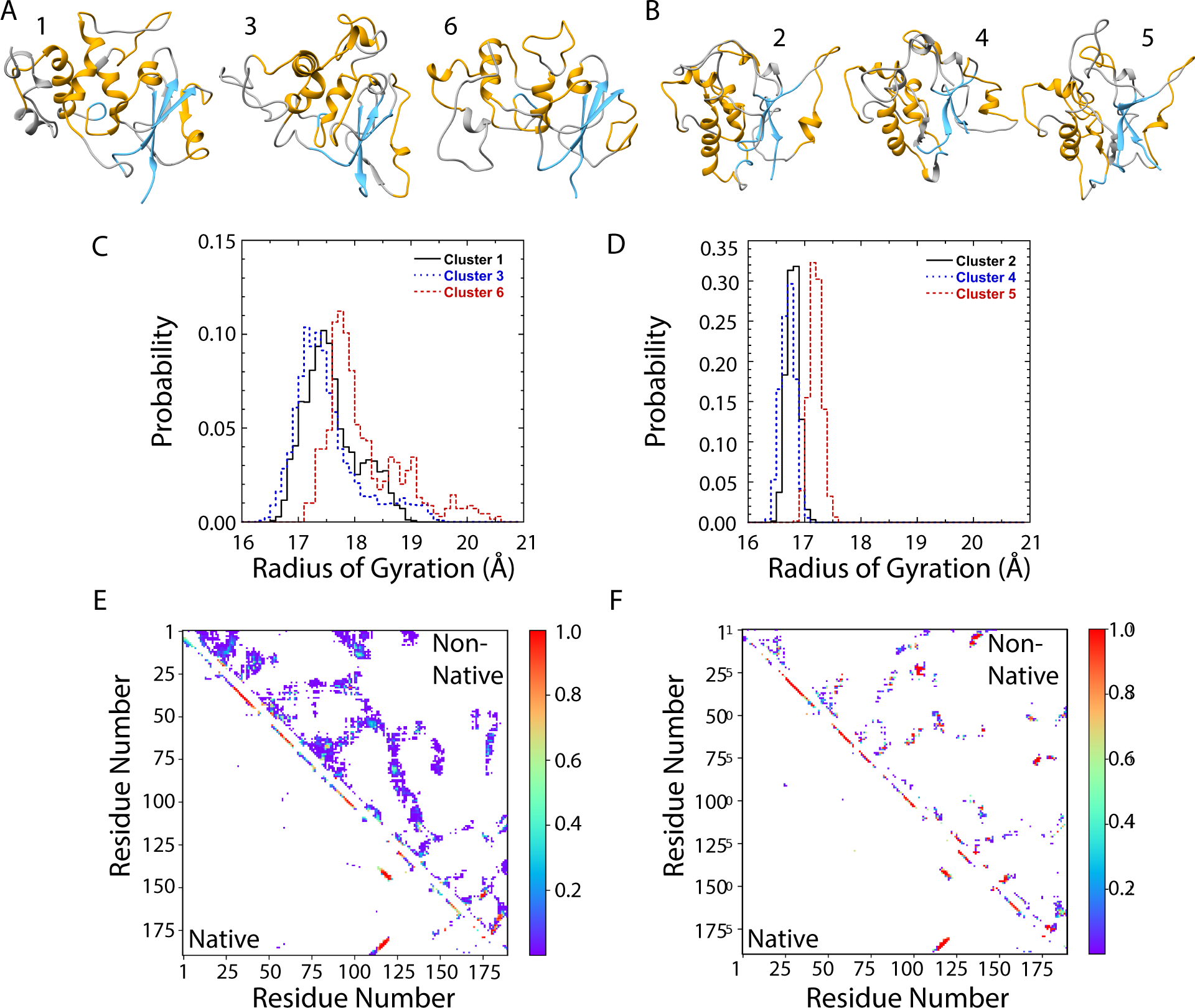
AAT190 conformational ensembles calculated by HDXer form two sub-populations with distinct properties. (A & B) Centroid structures for all six clusters calculated based on root mean square deviation (RMSD) labeled by cluster number, with regions that are α-helical, ý-strand or random coil in full-length, active AAT shown in gold, light blue and gray, respectively. Normalized radius of gyration (R_g_) distributions for (C) clusters 1 (black solid line), 3 (blue dotted line) and 6 (red dashed line), and (D) clusters 2 (black solid line), 4 (blue dotted line) and 5 (red dashed line). The change in y-axis scale between panels C and D emphasizes the relative compactness of clusters 2, 4 and 5. See Table S3 for statistical characterization of the Rg distributions. (E & F) AAT190 contact maps characteristic of each sub-population. Upweighted (E) cluster 1 and (F) cluster 2 showing that non-native contacts are longer lived and more localized in cluster 2 which is significantly more compact than is cluster 1. Native contacts present in the context of the full-length AAT structure are shown below the diagonal and non-native contacts are shown above the diagonal. The heat map indicates the relative occupancy determined using all frames for a given cluster. The number of frames and contact maps for each of the six clusters are in Table S3 and Figure S12, respectively.

## Discussion

Serpins are topologically complex and tend to form ordered polymers. Thus, while all proteins must avoid aggregation in order to fold efficiently (3), for serpins there is a particularly precarious balance between the flexibility required to incorporate long-range contacts and the sequestration of oligomerization prone elements. Our investigations of N-terminal AAT fragments suggest that, at least for AAT, this balance tips towards pathological polymerization only after the entire protein has been translated. Our biophysical investigations of the AAT190 structure along with recently reported data from Cabrita and coworkers on the AAT 1-191 N-terminal fragment (32) show that the N-terminal piece of the αβ subdomain, formed by almost half of the AAT sequence, is monomeric, relatively compact and contains labile, transient structure. AAT folding simulations (26, 27), proteolysis of ribosome-attached and free AAT nascent chains (32), and in-cell experiments on folding of the human inhibitory serpin antithrombin III (67) all suggest that tertiary structure consolidation in the labile N-terminal αβ subdomain is one of the last steps in folding of full-length serpins. The observed lability of this N-terminal subdomain is likely important because it provides the flexibility required to incorporate the AAT N-terminus into the full serpin fold as a late step in folding while also helping to avoid pathological oligomerization by both sequestering aggregation prone regions and providing a handle for the ER quality control machinery.

### From *in vitro* oligomerization propensity to in-cell clearance

For full-length serpins β-sheet expansion or β-strand mediated domain swaps and sheet expansion are hallmarks of serpin oligomerization/polymerization (14, 68–70). In the context of the full-length protein, the shortest N-terminal fragment, AAT190, contains six of the nine α-helices, a single, very short outer strand from sheet B and the strands 1 to 3 in sheet A (Fig. 1), resulting in two unsatisfied ý-strands, the very short strand 6B and strand 3A which the HDX-MS results show is relatively sequestered from solvent (Fig. 4 and 5). Lengthening the fragment to encompass the N-terminal subdomain of both the αβ and mainly β domains, AAT290, leaves all three β-sheets incomplete. The N-terminal mainly β subdomain, often referred to as the BC barrel, folds early both in experiments (25) and in simulations (26, 27); nonetheless, AAT290 contains at least four unsatisfied β-strands: 6B, 3A, 3B and 2C. Finally, AAT323 adds one additional dissatisfied strand, 6A. Thus, for the fragments, the observed increase in the probability of oligomerization and the number of monomers per oligomer correlates with an increase in the number of unsatisfied β-strands and suggests that the observed transition from monomeric AAT190 to progressively larger oligomers for AAT290 and AAT323 is β-strand mediated.

In the ER, where AAT and other secretory serpins fold and mature, protein fragments, misfolded proteins, and aberrant oligomers should be recognized by the quality control machinery and targeted for degradation. Failures in ER quality control can result in the accumulation of protein in the ER as observed for full-length secretory serpin mutants (71) and other proteins such as collagen (72, 73). We therefore asked whether wild-type AAT fragments or those containing the Siiyama (S53F) mutation would be successfully handled or would escape ER quality control.

The Siiyama mutation leads to the accumulation of full-length AAT polymers in the ER (33, 34, 74) and results in insoluble inclusion bodies when AAT190 containing the Siiyama mutation (AAT190S) is expressed in *E. coli*. While a slight increase in secretion for the longer fragments containing the Siiyama mutation (AAT290S and AAT323S) relative to the wild-type fragments was observed, in general even with the inclusion of the Siiyama mutation all of the fragments were efficiently recognized as misfolded and targeted for degradation (Fig. 3).

In the ER, three main interconnected quality control networks surveil maturing proteins: a classical chaperone network including Hsp40, Hsp70 and Hsp90 homologues, which recognizes exposed hydrophobic patches; glycan-mediated quality control comprising calnexin, calreticulin and enzymes that de- and re-glucosylate substrates (75); and thiol-dependent quality control (50, 76). Full-length AAT has three N-linked glycosylation sites, two in the αβ domain (N46 and N83) and one (N247) in the mainly β domain, as well a single cysteine, C232, in the mainly β domain (Fig. 1). N247 and C232 are missing in AAT190 but present in all other AAT constructs used for in-cell experiments (Table S2). In full-length human AAT all three N-linked glycosylation sites are usually 100% glycosylated (77, 78) and are known to mediate AAT binding to calnexin (79). For AAT fragments, N-linked glycosylation in CHO cells was confirmed by treatment with the glycosidase PNGase F, which resulted in the expected shift to lower molecular weight due to cleavage of the glycans (Fig. S13). Thus, all three quality control networks can interact with the AAT fragments and full-length AAT.

Hendershot and colleagues have shown that cross β-aggregation propensity as predicted by the Tango algorithm (80, 81) is correlated with recognition of aggregation-prone regions by the ER-resident chaperones Grp170, an HSP110 cognate, and two HSP40 family members, Erdj4 and Erdj5 (82). All three chaperones are part of the classical quality control network and can help target proteins for ERAD. Tango analysis of the AAT sequence yields high scores (>99/100) for strand 3A (residues 182-190), present in all of the fragments, and strand 4B (residues 370-376) near the C-terminus (Fig. S14), which is only present in full-length AAT. In addition, for the longer fragments, the middle of strand 3B (residues 248-255) has Tango scores in the 50s and is just C-terminal to Asn247, which should be glycosylated. In purified AAT190, the strand 3A region with high Tango scores (peptide 183-187) shows EX1 exchange (Fig. 4D) indicating concerted opening of this region, and such cooperative unfolding could facilitate chaperone recognition of this late folding region. It is therefore possible that transient exposure of aggregation-prone strands in the labile AAT fragments, which may be increased by lectin chaperone binding, facilitates efficient recognition of the fragments by the classical branch of ER quality control and subsequent degradation. Similar mechanisms might aid in the recognition and degradation of misfolded or slowly folding full-length AAT mutants. Interestingly, despite only moderate overall sequence conservation, in all fifteen inhibitory human secretory serpins, strands 3A and 4B show a high cross β-aggregation propensity as predicted by Tango (Fig. S15 and S16), suggesting that ER quality control could use similar mechanisms to surveil other secretory serpins.

In the longer AAT290 and AAT323 fragments, recognition by the ER quality control machinery must compete with oligomerization. But, in addition to the likely continued lability of structure in the N-terminal piece of the αβ domain, these longer residues also contain the single cysteine residue at 232, an additional N-linked glycosylation site at 247 and likely transient or long-lived exposure of hydrophobic residues in the BC barrel which should provide more recognition sites for ER quality control. For example, in AAT NHK, a truncated AAT mutant which behaves similarly to AAT323 in CHO cells (Fig. S5), C232 is important for recognition by the ER resident protein quality control factor α-mannosidase-like 1 protein (EDEM1) (83).

### Non-sequential folding and a key role for AAT190 lability

If the lability of the N-terminal αβ subdomain is important for surveillance of AAT folding by the ER quality control machinery, it would be useful to elucidate the AAT190 conformational ensemble. However, it is very difficult to do this in cells especially since AAT190 is quickly degraded by ERAD, and while proteasome inhibition by MG132 prolongs the lifetime of AAT190 in CHO cells, it also leads to the accumulation of detergent-insoluble AAT190 oligomers which would complicate experimental results (Fig. 3). Even in the absence of MG132, oligomerization is even more of a problem for the longer AAT290 and AAT 323 fragments. But, the intrinsic conformational ensemble of purified AAT190 should be similar to that in cells (32) allowing us to probe the properties of this ensemble using HDX-MS. HDX provides localized information on the solvent exposure of amide hydrogens. However, access of ER quality control machinery to a given sequence likely depends on both solvent exposure and the local structure of the protein chain. A detailed picture of the latter is not provided by HDX alone. By contrast, the HDXer-generated AAT190 conformational ensembles, which integrate HDX-MS peptide level data and MD simulations, provide information on both exposure and the local structural environment better representing what the ER quality control machinery actually encounters.

As expected from NMR characterization of the AAT 1 to 191 N-terminal fragment as a molten globule (32), AAT190 is extremely dynamic with most of the backbone amides exchanging within 10 sec (Fig. 4). The most protected regions in AAT190, helices B and E and strand 3A, are highly protected and slow to exchange in the full-length protein, highlighting the expected role of the protein sequence in protein structure formation and dynamics. As noted above, strand 3A has extremely high Tango scores and might be a particularly good recognition site for ER quality control. In AAT190 strand 3A is 60 percent exchanged within 1 min and the EX1 behavior suggesting concerted unfolding could allow molecular chaperones to easily access this region. The conformational ensembles generated by HDXer (Fig. 5 and S12) reiterate the importance of local secondary structure formation in AAT190, particularly in the β-strand region and show the ubiquity of non-native contacts. Many of the non-native contacts are close to native contacts and likely reflect slippage between native and non-native contacts. While AAT190 displays significant amounts of local secondary structure, the diversity of even the representative structures and transient nature of many of the contacts emphasize the predominantly disordered nature of this isolated serpin fragment. More globally, the results of the HDXer analysis show that this combination of HDX-MS data and simulations can be used to generate likely conformational ensembles even for proteins and protein fragments that are not well-structured.

Experimental studies (25, 32) and simulations (26, 27) all support AAT folding models in which docking of the N-terminal αβ subdomain to the body of the serpin and structural consolidation of this region is one of the final (if not the final) steps in serpin folding. Thus, native folding of the AAT N-terminal region requires the full-length protein and the observed varied conformational ensemble of the N-terminal subdomain (AAT190) may help nascent AAT chains avoid pathological polymerization even in the presence of polymerization promoting mutations such as Siiyama. Late folding and lability of an N-terminal piece of a non-sequential domain may not be limited to AAT. The 159-amino acid long *E. coli* protein dihydrofolate reductase (DHFR) consists of a non-sequential discontinuous loop domain (DLD) containing sequences from the N- and C-termini and a middle, sequential domain ABD (amino acids 38-106). HDX-MS studies of ribosome-attached nascent chains show that the N-terminal piece of the DLD exchanges quickly even for 126 amino acid long ribosome-attached nascent chains (84). For DHFR N-terminal lability appears to be mediated by the ribosome and is associated with faster folding of the full-length protein. Thus, for proteins with non-sequential domains, dynamic conformational heterogeneity of N-terminal subdomain and late structural consolidation, whether encoded in the protein sequence or mediated by interactions with the cellular quality control machinery including the ribosome, may aid in correct and efficient folding.

While it is tempting to speculate that lability and late folding of the N-terminal subdomain extends to serpins beyond AAT, it’s well known that members of the same protein family can have diverse folding mechanisms (85, 86), and even family members that populate similar folding intermediates can have different folding pathways (85). The functionally diverse serpin superfamily is quite large and is divided into 18 groups (also called clades) with groups A to I found in humans and other chordates (87). AAT, also known as serpinA1, is the most studied member of group A which includes not just protease inhibitors but also hormone transporters and angiotensinogen, peptides of which help control blood pressure. This diversity of function is reflected in the amino acid sequence identity of the 13 human group A serpins relative to AAT which ranges from 61 percent for SerpinA2 (AAT-related protein) to 21 percent for angiotensinogen with most group A serpins showing 30 to 40 percent identity to AAT. However, as found for AAT consolidation of structure in the N-terminal half of the serpin αβ domain is also a late step in the folding of human antithrombin III (SerpinC1) which shares 29 percent sequence identity with AAT (67). Therefore, our work and that of others suggest that structural lability, broad and dynamic conformational ensembles, and late folding of the N-terminal region is a general property of serpin folding, but further study of diverse serpins is required before generalizing the role of the N-terminal piece of the αβ domain in serpin folding.

### Experimental Procedures

Except as noted, reagents were purchased from Sigma Aldrich and Thermo Fisher Scientific and used without further purification.

#### Rapid Autonomous Fragment Test (RAFT) prediction of AFUs

The RAFT score, *R(S)*, for any given sequential protein segment S of length *l* is calculated as the difference between the number of internal-internal (*II)* and the number of internal-external (*IE*) contacts (43):

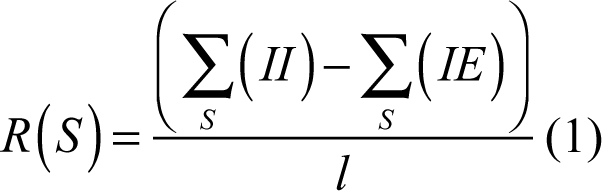

High scoring segments have many more internal-internal contacts and are considered more likely to fold autonomously. For AAT the top scoring sequential structures and associated RAFT scores are shown in Figure S2.

#### AAT Constructs

For expression in *E. coli* the mature (i.e, no signal sequence) AAT reference sequence M1V (UniProt P01009 (44)) with the addition of the Cys232Ser (C232S) mutation was cloned from the pcDNA plasmid into the pQE-30 plasmid (Qiagen) resulting in a sequence with an N-terminal 6XHis tag and TEV cleavage site (see Table S4 for primer sequences). Site-directed mutagenesis was used to introduce a stop codon at positions 191, 291 and 324 in the AAT sequence for fragments 1-190, 1-290 and 1-323, respectively and Sanger sequencing (Genewiz, Inc.) was used to confirm all sequences (see Table S1 for amino acid sequences).

Mature, full-length AAT M1V containing the C232S mutation was expressed in *E. coli* using the pEAT8-137 plasmid (88, 89). The original pEAT8-137 plasmid encodes the AAT wild-type variant M2 with two mutations (R101H and E376D) relative to the more common M1V sequence (90). These mutations were reverted and confirmed by sequencing (Genewiz, Inc) resulting in the pEAT8-137 plasmid encoding M1V with the C232S mutation. See Tables S4 and S5 for DNA primers and the amino acid sequence, respectively.

For expression in CHO cells, the N-terminus of all constructs consisted of the AAT signal sequence for proper targeting to the ER, the first 3 residues (Glu Asp Pro) of the mature sequence to ensure signal sequence cleavage, a Myc tag for immunoblotting followed by 6XHis and TEV recognition sequences to mimic the N-terminus used for *E. coli* expression. The required AAT M1V constructs were produced by amplifying the tag and linker sequence from the pcDNA3.1 AAT plasmid encoding C-terminally tagged AAT M1V and inserting this fragment between P3 and Q4 in pcDNA3.1 using Gibson assembly and inserting stop codons as needed for each fragment and for full-length AAT M1V (see Table S4 for DNA primer sequences). All sequences were confirmed by Sanger sequencing (Genewiz). See Table S2 for amino acid sequences.

#### AAT construct purification

Fragments were expressed and purified using a protocol similar to that published by Dolmer and Gettins for a stabilized version of AAT323 (21). *E. coli* strain BL21(DE3) cells transformed with the pQE-30 plasmid containing the desired fragment were used to inoculate 1 liter of Luria Broth (LB) containing 100 μg/ml ampicillin. Cultures were grown at 37 °C and 250 rpm until an OD_600_ of 0.6 to 0.7 was reached at which point protein expression was induced using 1 mM **i**sopropyl β-D-thiogalactoside (IPTG) and the cultures were then grown for a further 4 hr. *E. coli* cells were harvested by centrifugation, resuspended in 50 mM potassium phosphate, 150 mM NaCl pH 7.4 (KP150) containing protease inhibitors (1 mM PMSF and 1 μg/ml pepstatin A), flash frozen in liquid nitrogen and stored at -80 °C. Cells were defrosted and lysed using a microfluidizer (Microfluidics model 110M) or sonication, and inclusion bodies containing the fragments were isolated by centrifugation and washed three times. Inclusion bodies were then solubilized in KP150 containing 6 M urea (unfolding buffer) for 1 hr at room temperature, and applied to a Ni-NTA agarose column (Qiagen) equilibrated with unfolding buffer. The column was washed with unfolding buffer plus 25 mM imidazole and AAT fragments were eluted with unfolding buffer plus 500 mM imidazole. The fragments were refolded dropwise at 4 °C using gentle stirring and a 1:50 dilution into KP150. The folded fragments were concentrated using a Ni-NTA column and eluted in 50 mM potassium phosphate, 300 mM NaCl pH 7.4 (KP300) plus 500 mM imidazole. Purified fragments were dialyzed into KP150 containing 1mM DTT and 1 mM EDTA (cleavage buffer).

TEV protease was expressed and purified as described previously (91), and 4.6 μM of 6XHis tagged TEV protease was incubated with AAT fragments in KP150 at room temperature for 1 hr followed by an incubation at 4 °C for 16 hr. The AAT fragment/TEV solution was applied to a Ni-NTA column. The column was washed with 50 mM potassium phosphate, 150 mM NaCl, 25 mM imidazole, pH 7.4, and the cleaved fragment eluted at a higher imidazole concentration (50 mM). The cleaved fragments were then dialyzed against KP150 and stored at 4 °C at concentrations ranging from 8 to 10 µM. (Attempts to freeze the fragments at -20 or -80 °C resulted in aggregation.)

#### Full-length AAT purification

Full-length AAT M1V containing the C232S mutation in the plasmid pEAT8-137 was expressed as a soluble protein in *E. coli* BL21(DE3) cells and purified using two anion exchange columns as described previously (20, 92). The inhibitory activity was assayed against bovine trypsin as described (92). Purified full-length AAT was active with a stoichiometry of inhibition of 1.0 to 1.1 for trypsin.

#### Protein Concentrations

Protein concentrations were calculated from the absorbance at 280 nm. For full-length AAT, the published extinction coefficient at 280 nm is 23,484 M^-1^ cm^-1^ (0.53 ml/(mg*cm)) (93). Fragment extinction coefficients were calculated using the amino acid sequence and the ExPasy ProtParam server (94) and are listed in Table S1.

#### AUC sedimentation velocity (SV) experiments

Purified proteins were dialyzed into 50 mM potassium phosphate, pH 7.4 for 16 hr at 4 °C and diluted to obtain the desired protein concentration. SV experiments were performed using a Beckman Coulter ProteomeLab XL-I or Beckman Coulter Optima XL-A analytical ultracentrifuge equipped with an optical detection system. All runs were performed in a double sector charcoal filled epon centerpieces with quartz windows. The volume for each sample was 400 μL and 410 μL of buffer used in the reference sector so that the meniscus of the reference was offset and did not obscure the sample meniscus. The runs were carried out in an An60Ti rotor at either 42,000 or 55,000 rpm, 20°C, and the absorbance data (1.2 cm path length) were acquired at 230 nm for AAT190, which contains no Trp residues, or 280 nm for longer fragments which contain two Trp residues. For AAT290 and AAT323 SV experiments were performed for two biological replicates. For AAT190 SV experiments were performed for four biological replicates.

The acquired absorbance data were fit to the continuous coefficient distribution (c(s)) model using SEDFIT (95). The buffer density (ρ), 1.0051 g/ml, and buffer viscosity (11), 0.01018 Poise, at 20 °C were calculated for 50 mM potassium phosphate pH 7.4 using SEDNTERP (96) and the default value, 0.73 ml/g, was used for the macromolecular partial specific volume, 𝜈. The distribution obtained from the SV data was discretized with a grid of 150 s values between 0 and 12 S for AAT190 and 1 to 20 S for AAT290 and AAT323. For diffusion deconvolution, a scaling law for compact particles was used with the average frictional ratio allowed to change in the fits, along with the initial meniscus position. The radial independent (RI) and time independent (TI) noise were accounted for in the SV data analysis using algebraic noise decomposition based on least squares modeling of the sedimentation data (97). The best-fit c(s) model was based on the minimization of the root-mean-square deviation of the fit and on the shape of the residuals plot and bitmap. The values of the sedimentation coefficient at 20 °C in water (*s_20,w_*) were obtained by integration of the fitted c(s) distributions. The 68% confidence intervals associated with *s_20,w_* were determined by F-statistics considering variations in the meniscus position over the quality of the best-fit c(s) distribution in SEDFIT.

#### Cell Culture

CHO-K1 (lot 62960170) cells were purchased from ATCC. Cells were checked for mycoplasma contamination using a universal mycoplasma detection kit (catalog no. 30-012K, ATCC). Cells were grown in α−minimal essential medium supplemented with 10% FEBS and 1% penicillin/streptomycin at 37 °C and 5% CO_2_. Cells in 3.5 cm dishes were transfected with 1 μg pcDNA3.1 plasmid using polyethylenimine (PEI) and Opti-MEM reduced serum media and incubated for 20 hr at 37 °C and 5% CO_2_. Control samples were grown for a further 20 hr. For proteasome inhibition experiments, after 20 hr of incubation MG132 was added to the media to a final concentration of 20 μM and the cells were incubated for an additional 20 hr prior to collecting samples.

#### SDS-PAGE and Western blots

Cell media was collected and centrifuged at 14000 x g for 5 min at 4 °C and the clarified cell media was collected. Cells were washed with cold PBS and lysed in lysis buffer (20mM MES, 100mM NaCl, 30mM Tris-HCl pH 7.5, 0.5% Triton X-100) supplemented with protease inhibitors (20 mM *N*-ethylmaleimide, 400 μM phenylmethylsulfonyl fluoride, 50 μM calpain inhibitor 1, 1μM pepstatin, 10 μg/μl aprotinin, 10 μg/μl leupeptin). Following lysis, cells were vortexed at high speed for 5 min at 4 °C and centrifuged at 14000 x g for 5 min at 4 °C. Post-nuclear supernatants were collected. To prepare the Triton X-100 (detergent) insoluble sample, the pellet from the cell lysate centrifugation was washed with lysis buffer and resuspended in 100 mM Tris pH 8, 1% SDS by vortexing at high speed for 5 min at room temperature, incubation at 95 °C for 10 min, and (following a 10:1 dilution with lysis buffer) sonication for 30 sec at 4 °C. The detergent-insoluble sample was further diluted 5:1 in lysis buffer.

Proteins in the media, cell lysate and detergent-insoluble samples were precipitated by vortexing with 10% v/v trichloroacetic acid (TCA). Pellets obtained from 5 min of fast centrifugation were washed with acetone, centrifuged for a further 5 min, isolated and dissolved in reducing sample buffer (9% SDS,15% glycerol, 30mM Tris pH 6.8, 0.05% bromophenol blue) by high speed vortexing at room temperature for 5 min and incubation at 95 °C for 5 min. Samples were centrifuged at 14000 x g for 5 min and 10 μl of each sample was loaded on a 9% SDS-PAGE gel.

Following SDS-PAGE, gels were washed with ultrapure water, and transfer membranes (Immobilon-FL, Millipore) were pre-treated according to the manufacturer’s instructions. Proteins were transferred to the membrane using a semi-dry transfer unit (Hoefer TE77XP), and blots were blocked in 5% milk, 2% BSA in PBS for 1 hr under gentle shaking at room temperature. A 1:1000 dilution of the primary monoclonal mouse Myc tag 9B11 antibody (Cell Signaling, lot 24) in 5% milk, 2% BSA, PBST (2.7 mM KCl, 1.5 mM KH_2_PO_4_, 136.9 mM NaCl, 8 mM Na_2_PHO_4_, 0.5% Tween 20, pH 7.4) was added to the blots and blots were incubated overnight at 4 °C under gentle shaking. Blots were then washed with PBST and incubated with IRDye 800CW-conjugated goat anti-mouse secondary antibody (LI-COR Biosciences, lot C40528-02) in a 5% milk, 2% BSA, 0.02% SDS, PBST solution for 1 hr at 25 °C under gentle shaking. Blots were then washed with PBST and imaged using a LI-COR Odyssey CLx imaging system

#### HDX-MS

The coverage maps for full-length AAT M1V and AAT190 were obtained from undeuterated 8 μM samples in 50 mM potassium phosphate pH 7.4. Samples were diluted 1:5 with 100 mM potassium phosphate, 1 M guanidinium chloride (GdmCl), pH 2.5 at 0 °C. The sample was immediately injected into a Waters HDX nano Acquity ultra-performance liquid chromatography (UPLC) system (Waters, Milford, MA) with in-line pepsin digestion (Waters Enzymate BEH column). Peptic fragments were trapped on an Acquity UPLC BEH 150 C18 column peptide trap and separated on an Acquity UPLC HSS T3 C18 attached to an Acquity UPLC BEH C18 guard column. A 7 min, 5 to 35% (vol/vol) acetonitrile (0.1% formic acid) gradient was used to elute peptides directly into the Synapt G2 mass spectrometer (Waters). MS data were acquired with a 20- to 30-V ramp trap collision energy for high-energy acquisition of product ions and lock mass (Leu-Enk) for mass accuracy correction. Peptides were identified using the Protein Lynx Global Server 2.5.1 (PLGS) from Waters.

For HDX experiments 8 μM protein in 50 mM potassium phosphate pH 7.4 was incubated at 25 °C for 5 min and then diluted 1:5 into deuteration buffer (50 mM potassium phosphate, 150 mM NaCl, pD 7.4) over a range of time points (0 sec, 10 sec, 1 min, 10 min, 1 hr and 2 hr). Exchange was quenched by a 1:1 dilution into 100 mM potassium phosphate, 1M GdmCl, pH 2.5 at 0 °C.

All deuteration reactions were performed in triplicate. To account for back exchange, where the deuterium exchanges for hydrogen following the quench, a fully deuterated control sample was generated by incubating 8 to 10 μM protein in deuteration buffer containing 6 M deuterated GdmCl for 2 hr at 25 °C. Deuterium uptake by the identified peptides, in the time dependent samples and fully deuterated control, was determined using the DynamX 3.0 software (Waters). For peptides displaying bimodal spectra, the two peaks were de-convoluted using HX-Express v2 (98, 99). HDX-MS experiments were repeated for two biological replicates of AAT190. HDX-MS data quality metrics are in Table S6.

Prior to all MS experiments, AAT190 samples were checked for oligomerization using AUC and samples that were at least 80% monomeric were subjected to HDX-MS. Oligomers could lead to multiple peaks in the HDX-MS spectra. However, at 10 sec none of the AAT190 peptides show more than one peak, suggesting that oligomers do not make a significant contribution to the observed spectra. This lack could arise because the oligomers are in rapid equilibrium with monomers, due to heterogeneities in the oligomer population or because integrating the c(s) distribution for all sedimentation values beyond the monomer peak (>3 S) overestimates the oligomer population.

#### Conventional MD Simulations

For AAT190, the structure of residues 23 to 190 was excised from the longer structure (PDB: 1QLP (36)). The CHARMM GUI (37) was used to model in the missing 22 N-terminal residues for both AAT190 and the full-length structure. Input files for OpenMM (100) were generated using the CHARMM-GUI Glycan Reader & Modeler web application (101) and the CHARMM36m additive force field (37) with TIP3P water (102). The proteins were solvated in a periodic water box containing 0.15M KCl, with box boundaries buffering 1 nm away from the molecule. A force switching function was applied from 1.0 to 1.2 nm for the Lennard-Jones interaction calculations and the particle-mesh Ewald method (103) (Ewald error tolerance of 0.0005) was used to calculate long-range electrostatic interactions. Before the simulation run, the system energy was minimized using the limited memory Broyden-Fletcher-Goldfarb-Shanno (L-BGFS) method and then equilibrated for 125 psec using a 1 fsec integration step. For both equilibration and minimization, positional restraints were applied to the protein backbone and side chains using a force constant of 400 and 40 kJ/mol/A^2^, respectively. During production, a 2 fsec time step was used for integration with temperature and pressure held constant at 298.15 K and 1 bar, respectively. Pressure was isotopically held constant at 1 bar using a Monte Carlo barostat with a pressure coupling frequency of 2 psec. The resulting equilibrated and energy minimized input was simulated for 1 µsec, with structural coordinates written to the trajectory every 20 psec of simulated time, resulting in a total of 50,000 frames. Results were visualized and analyzed using VMD (104).

#### Enhanced Sampling Using Simulated Tempering (ST)

In tempering methods, elevated temperatures are used to help the protein chain escape from local minima. Thus, to better explore the conformational landscape, the ST enhanced sampling method (105–108) was applied to AAT190. ST simulations were performed using the AAT190 structure, CHARMM36m force field (109), OpenMM (100) and the minimization and equilibration procedures described above for the conventional MD simulations. To ensure thorough exploration of the conformational landscape, four 5.0 X 10^7^ step long (1 fsec/step) ST runs were performed using 100 possible equally spaced transition temperatures between the minimum temperature of 300K and maximum temperatures of 400K, 440K, 480K and 500K, respectively. Each temperature was assigned a weight to promote equal sampling at the given temperatures throughout the course of the simulation (107) and ST was performed with temperature transitions every 25 psec. The resulting four trajectories containing 50,000 frames (20 psec/frame) were then combined for the HDXer analysis.

#### Generating Reweighted Conformational Ensembles Using HDXer

The HDXer approach developed by Bradshaw, Lee, Forrest and co-workers uses HDX-MS experimental data, enhanced sampling MD simulations and maximum entropy methods to reweight conformational ensembles to better fit the experimental results (39, 60). In this approach, the protection factor for residue i, *P_i_*, is calculated from the simulated trajectories using the phenomenological equation from Best and Verndruscolo (110):

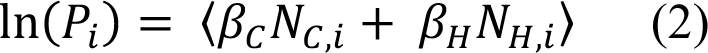

Where <> indicates the ensemble average over all 200,000 frames in the consolidated ST trajectories, *N_c,I_* is the number of non-hydrogen atoms within 6.5 Å of the backbone nitrogen atom of the residue omitting the two neighboring residues, *N_H,i_* is the number of hydrogen bonds involving the backbone amide of residue *i*, and *ý_c_* = 0.35 and *ý_H_* = 2.0 are the suggested scaling factors. These protection factors were then used to calculate the time dependence of the peptide level deuterium uptake, 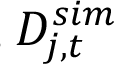 for all experimentally observed peptides exhibiting EX2 exchange:

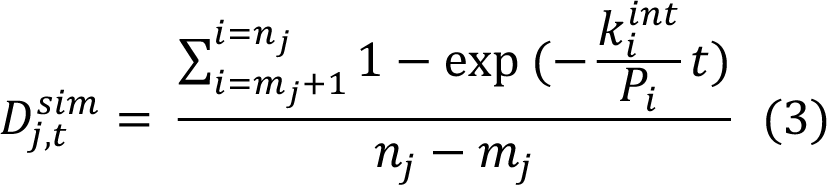

where *t* is time, *n_j_* and *m_j_* are the starting and ending residues for peptide j and *k_int_^i^* is the intrinsic rate constant for exchange (111, 112). These calculated 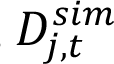 values and maximum entropy reweighting are then used to fit the experimental HDX-MS data and to reassign weights for each frame in the 200,000 frame combined ST trajectories as described in detail by Bradshaw and co-workers (39). Initial weights, 𝛺_*i*_, for each frame are simply equal to 1/n where n is the total number of frames. After reweighting, an optimized weight (Ω _-_) for each frame is obtained such that frames that more closely represent the data are given Ω _*f*_ > Ω _*i*_, while frames that do not represent the data well are given values of Ω _*f*_ < Ω _*i*_.

To prevent overfitting, the tightness of fit between the simulated and experimental data can be controlled by the γ parameter. The choice of γ affects both the final mean squared deviation (MSD) of the fitted data to the experimental data and the apparent work (*W*_app_, in kJ/mol) applied to the initial ensemble to adjust the conformational populations. The point of inflection in the decision plot of *W*_app_ *vs.* MSD (Fig. S11) denotes the onset of overfitting, characterized as a rapid increase in *W*_app_ for very little reduction in MSD. For AAT190 this rapid increase in *W_app_* is observed at 0.439 kJ/mol yielding a γ value of 2.

#### Dimensional Reduction using Time Lagged Independent Component Analysis (tICA)

The output of the HDXer method is a large number of structures (frames) that can be difficult to interpret and clustering methods such as tICA (61) can be used to identify similarities between structures facilitating structural interpretations of the HDX-MS data. TICA clustering was performed in PyEmma (62) using backbone dihedral angles as the input. The number of clusters and cluster boundaries were determined and the reassigned HDXer weights, Ω _-_, were mapped to the tICA clusters. Iterative tICA clustering, preferred for complex ensembles such as those resulting from ST and other enhanced sampling techniques, was performed by isolating individual clusters and subjecting each to further tICA based clustering. Iterative clustering was terminated when clusters contained <1000 structures and the RMSD for all structures in the cluster was <3 Å. To identify clusters containing frames upweighted by HDXer the average weight reassignment for a given cluster was then calculated as the mean of the log fold change in weight, log(Θ_f_/Θ_i_), with upweighted clusters displaying a mean log(Θ_f_/Θ_i_) > 0. Z scores were calculated to determine the statistical significance of the change in weights for individual upweighted clusters:

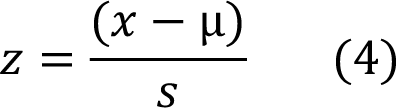

where 𝑥 is the mean log(Θ_f_/Θ_i_) of a given cluster, µ is the mean log(Θ_f_/Θ_i_) for all clusters and s is the standard deviation in average log(Θ_f_/Θ_i_) values for all clusters. Z scores were then converted to percentile using z score to percentile table (a z score > 1.7 is a significance ≥ 95%), from which a p-value was derived. Clusters with p-values <0.05 were selected as significantly upweighted clusters and were ranked based on log(Θ_f_/Θ_i_) value. TICA clustering identified six out of 186 total clusters as upweighted by HDXer. For comparison to the results of AUC SV experiments, theoretical values of the frictional ratio were calculated for the computational ensembles using the compiled trajectory for each of the six clusters as inputs to the HullRad webserver (Fig. S11B) (63).

To characterize the conformational landscape of the resulting AAT190 ST clusters, kernel density estimation plots (KDE plots) were generated as a function of hydrodynamic radius and the distance between residue F61 in helix B and L184 in s3A two conserved shutter residues that pack against each other in the folded full-length protein (113) (Fig. S11C). Distance calculations were performed for each frame of the upweighted clusters as well as the full-length AAT MD simulation by measuring the distance between the α-carbons using MDTraj (114). The representative structures and R_g_ distributions shown in Figure 5 were also calculated using MDTraj (114). The secondary structure for the representative structures in Figures 5A and 5B was assigned by UCSF Chimera (115) using the Kabsch and Sander method (116).

#### Contact Occupancy

For contact occupancy analysis of the full-length AAT structure (Fig. 1 and S1) and the clusters from the HDXer analysis (Fig. 5 and S12), contacts were defined between alpha carbons using a cut-off distance of 7.5 Å. Distances between α carbons were calculated with MDTraj (114) and contacts for each frame were identified using a Python script which also calculated the occupancy; the percentage of frames in which each contact was formed. For comparison of native versus non-native contacts, native contacts were defined as contacts between residues 1-190 in the context of the full-length AAT simulation. To eliminate extremely transient interactions, a contact was defined as native only if it is formed in ≥ 10% of the frames. For the non-native contact maps of the clusters, all native contacts defined from the full-length AAT simulation were eliminated from the map, leaving only the occupancy of non-native contacts.

### Predicting β Aggregation Propensities

The mature amino acid sequences (no signal sequence or pro sequence) of human extracellular inhibitory serpins and squamous cell carcinoma antigen 1 (SCCA1, SerpinB3), a human intracellular inhibitory serpin, were retrieved from the Uniprot database (44). The Tango server (http://tango.crg.es/) (80, 81) was used to predict cross β aggregation propensity using parameters selected to mimic the ER: unprotected N and C termini, 310.15K (37 °C) temperature, pH 7.2 and 175 mM ionic strength. The sequence parameter was set to 1 μM but note that this parameter has no effect on the calculation. The serpin sequences were aligned using Clustal Omega (117) and the alignment was displayed using ESPript 3.0 (118). Alignments and Tango cross β aggregation scores for the s3A and s4B regions are shown in Figures S15 and S16, respectively.

## Data Availability Statement

All data are available from the authors upon request. The pEAT8-137 plasmid for expressing full-length AAT M1V is available from Addgene (https://www.addgene.org/173807). Please send requests for other reagents or data to Anne Gershenson or Daniel Deredge

**Author Contributions:** A.G., D.D., D.N.H., L.M.G., P.L.W. & W.K. conceptualization; A.G., D.D., D.N.H., H.K., K.C.K., L.M.G., P. L.W., U.K. & W.K. methodology; D.D., H.K., K.C.K., P.L.W., U.K. & W.K. investigation; A.G., D.D., D.N.H., H.K., K.C.K., L.M.G., P.L.W., U K. & W.K. formal analysis; A.G., H.K., K.C.K., P.L.W. & U.K. writing-original draft; A.G., D.D., D.N.H., H.K., K.C.K., L.M.G., & P.L.W. writing-review & editing; A.G., D.D., D.N.H., L.M.G. & P.L.W. supervision; A.G. project administration.

## Acknowledgements

**Acknowledgements:** We thank Prof. Eugenia M. Clerico, Dr. Beena Krishnan, Dr. Santosh M. Kumar and a number of University of Massachusetts Amherst undergraduates including Austin Hachey, Matthew Lee and Lauren Prentis for their contributions to this research. We also thank Dr. Kael F. Fischer for running the RAFT calculations and Lizz Bartlett, the former director of the University of Massachusetts Amherst Biophysical Characterization core facility, for assistance with AUC data collection and analysis.

**Funding and Additional Information:** This work was supported by the Alpha-1 Foundation (D.N.H., L.M.G., A.G. and P.L.W.); NIH/NIGMS awards R01 GM128985 (A.G. subcontract), R01 GM086874 (D.N.H.) and R35 GM118161 (L.M.G.). This research was conducted while Anne Gershenson and Weiwei Kuo were employed at the University of Massachusetts Amherst. The opinions expressed in this article are the authors’ own and do not reflect the view of the National Institutes of Health, the Food and Drug Administration, the Department of Health and Human Services, or the United States government.

**Conflicts of Interest:** The authors declare that they have no conflicts of interest with the contents of this article.

## Abbreviations

*–* The abbreviations used are:

AAT: α_1_-antitrypsin
AAT190: 1-190 AAT fragment
AAT290: 1-290 AAT fragment
AAT323: 1-323 AAT fragment
AFU: autonomous folding unit
AUC: analytical ultracentrifugation
CD: far UV circular dichroism
CHO: Chinese hamster ovary
ER: endoplasmic reticulum
ERAD: ER-associated protein degradation
HDX-MS: hydrogen-deuterium exchange mass spectrometry
LC: liquid chromatography
MSD: mean squared deviation
RCL: reactive center loop
ST: simulated tempering
SV: sedimentation velocity
TCA: trichloroacetic acid
tICA: time lagged independent component analysis

## Notes

### Competing Interest Statement

The authors have declared no competing interest.

