## Supporting Information for "The conformational landscape of a serpin N-terminal subdomain facilitates folding and in-cell quality control"

### Supplementary Experimental Procedures

#### [$^{35}$S]-Met/Cys Metabolic Labeling and Pulse-Chase

Twenty hr post-transfection the cell media was removed from the 3.5 cm dishes and replaced with modified Dulbecco's eagle medium (DMEM) which lacked sodium pyruvate, L-Gln, L-Cys and L-Met and contained 4.5 g/L D-glucose and 55 μCi of EasyTag Express $^{35}$S Protein Labeling Mix [$^{35}$S]-Met/Cys ([$^{35}$S]-Met :[$^{35}$S]-Cys: = 73:22) (PerkinElmer Life Sciences). The Chinese hamster ovary (CHO) cells were radiolabeled for 30 min. Immediately after the pulse, cells were washed with PBS and chased for the indicated time using regular growth media. At each timepoint, cells were washed with cold PBS and lysed in lysis buffer (20 mM MES, 100 mM NaCl, 30 mM Tris-HCl pH 7.5, 0.5% Triton-X100) supplemented with protease inhibitors (20 mM *N*-ethylmaleimide, 400 μM phenylmethylsulfonyl fluoride, 50 μM calpain inhibitor 1, 1 μM pepstatin, 10 μg/μl aprotinin, 10 μg/μl leupeptin). Media and detergent (Triton X-100)-insoluble fractions were collected where indicated (Fig. S5). After the pulse-chase, AAT was immunoprecipitated from cell lysate, media and Triton X-100 insoluble cellular fractions using polyclonal rabbit anti-human AAT antibody (Agilent Dako, catalog number A 0012, lot 20028622).

### Supplementary Tables and Figures

**Table S1.** Protein sequences for AAT fragments expressed in *E. coli* with the N-terminal 6XHis tag in vermillion, the TEV protease recognition sequence in blue, the TEV cleavage site indicated by "/" and, for the longer constructs, the Cys232Ser mutation is purple.

| | | After TEV Cleavage | |
|---|---|---|---|
| **AAT Construct** | **Amino Acid Sequence**[1] | **Molecular Weight (kDa)**[2] | **Calculated $\varepsilon_{280}$ ($M^{-1}$ $cm^{-1}$)**[2] |
| 1-190 (AAT190) | -17 MRGSHHHHHH GENLYFQ/<br>1 GDPQGDAAQK TDTSHHDQDH PTFNKITPNL<br>31 AEFAFSLYRQ LAHQSNSTNI FFSPVSIATA<br>61 FAMLSLGTKA DTHDEILEGL NFNLTEIPEA<br>91 QIHEGFQELL RTLNQPDSQL QLTTGNGLFL<br>121 SEGLKLVDKF LEDVKKLYHS EAFTVNFGDT<br>151 EEAKKQINDY VEKGTQGKIV DLVKELDRDT<br>181 VFALVNYIFF | 21,325 | 5,960 |
| 1-290 C232S (AAT290) | -17 MRGSHHHHHH GENLYFQ/<br>1 GDPQGDAAQK TDTSHHDQDH PTFNKITPNL<br>31 AEFAFSLYRQ LAHQSNSTNI FFSPVSIATA<br>61 FAMLSLGTKA DTHDEILEGL NFNLTEIPEA<br>91 QIHEGFQELL RTLNQPDSQL QLTTGNGLFL<br>121 SEGLKLVDKF LEDVKKLYHS EAFTVNFGDT<br>151 EEAKKQINDY VEKGTQGKIV DLVKELDRDT<br>181 VFALVNYIFF KGKWERPFEV KDTEEEDFHV<br>211 DQVTTVKVPM MKRLGMFNIQ HSKKLSSWVL<br>241 LMKYLGNATA IFFLPDEGKL QHLENELTHD<br>271 IITKFLENED RRSASLHLPK | 33,089 | 18,450 |
| 1-323 C232S (AAT323) | -17 MRGSHHHHHH GENLYFQ/<br>1 GDPQGDAAQK TDTSHHDQDH PTFNKITPNL<br>31 AEFAFSLYRQ LAHQSNSTNI FFSPVSIATA<br>61 FAMLSLGTKA DTHDEILEGL NFNLTEIPEA<br>91 QIHEGFQELL RTLNQPDSQL QLTTGNGLFL<br>121 SEGLKLVDKF LEDVKKLYHS EAFTVNFGDT<br>151 EEAKKQINDY VEKGTQGKIV DLVKELDRDT<br>181 VFALVNYIFF KGKWERPFEV KDTEEEDFHV<br>211 DQVTTVKVPM MKRLGMFNIQ HSKKLSSWVL<br>241 LMKYLGNATA IFFLPDEGKL QHLENELTHD<br>271 IITKFLENED RRSASLHLPK LSITGTYDLK<br>301 SVLGQLGITK VFSNGADLSG VTE | 36,456 | 19,940 |

[1]For the AAT fragments, the N-terminus following TEV protease cleavage is numbered as 1. The TEV recognition sequence results in the glutamic acid to glycine mutation at the first residue (E1G).

[2]Calculated using Expasy ProtParam (https://web.expasy.org/protparam/) (1).

**Table S2.** Protein sequences for AAT fragments and full-length proteins expressed in CHO cells with the signal sequence in bluish green (negative numbers); the mature AAT sequence in black (positive numbers); the N-terminal myc linker and 6XHis tag in vermillion; the TEV protease recognition sequence in blue; the locations of N-linked glycans in red; serine 53, the residue mutated to phenylalanine in the Siiyama variant, is highlighted in yellow; and for longer fragments the single cysteine residue is in purple.[1] For the *null* Hong Kong (NHK) variants amino acids after the frameshift are in pink.

| AAT Construct | Amino Acid Sequence |
|---|---|
| **1-190 (AAT190)** | 123<br>-24 MPSSVSWGIL LLAGLCCLVP VSLAEDPEQK LISEEDLNSA VDHHHHHHEN<br>4<br>LYFQGQGDAA QKTDTSHHDQ DHPTFNKITP NLAEFAFSLY RQLAHQS**N**ST<br>49 NIFF**S**PVSIA TAFAMLSLGT KADTHDEILE GLNF**N**LTEIP EAQIHEGFQE<br>99 LLRTLNQPDS QLQLTTGNGL FLSEGLKLVD KFLEDVKKLY HSEAFTVNFG<br>149 DTEEAKKQIN DYVEKGTQGK IVDLVKELDR DTVFALVNYI FF |
| **1-290 (AAT290)** | 123<br>-24 MPSSVSWGIL LLAGLCCLVP VSLAEDPEQK LISEEDLNSA VDHHHHHHEN<br>4<br>LYFQGQGDAA QKTDTSHHDQ DHPTFNKITP NLAEFAFSLY RQLAHQS**N**ST<br>49 NIFF**S**PVSIA TAFAMLSLGT KADTHDEILE GLNF**N**LTEIP EAQIHEGFQE<br>99 LLRTLNQPDS QLQLTTGNGL FLSEGLKLVD KFLEDVKKLY HSEAFTVNFG<br>149 DTEEAKKQIN DYVEKGTQGK IVDLVKELDR DTVFALVNYI FFKGKWERPF<br>199 EVKDTEEEDF HVDQVTTVKV PMMKRLGMFN IQH**C**KKLSSW VLLMKYLG**N**A<br>249 TAIFFLPDEG KLQHLENELT HDIITKFLEN EDRRSASLHL PK |
| **1-323 (AAT323)** | 123<br>-24 MPSSVSWGIL LLAGLCCLVP VSLAEDPEQK LISEEDLNSA VDHHHHHHEN<br>4<br>LYFQGQGDAA QKTDTSHHDQ DHPTFNKITP NLAEFAFSLY RQLAHQS**N**ST<br>49 NIFF**S**PVSIA TAFAMLSLGT KADTHDEILE GLNF**N**LTEIP EAQIHEGFQE<br>99 LLRTLNQPDS QLQLTTGNGL FLSEGLKLVD KFLEDVKKLY HSEAFTVNFG<br>149 DTEEAKKQIN DYVEKGTQGK IVDLVKELDR DTVFALVNYI FFKGKWERPF<br>199 EVKDTEEEDF HVDQVTTVKV PMMKRLGMFN IQH**C**KKLSSW VLLMKYLG**N**A<br>249 TAIFFLPDEG KLQHLENELT HDIITKFLEN EDRRSASLHL PKLSITGTYD<br>299 LKSVLGQLGI TKVFSNGADL SGVTE |
| ***Null* Hong Kong (NHK)[2]** | 123<br>-24 MPSSVSWGIL LLAGLCCLVP VSLAEDPQGD AAQKTDTSHH DQDHPTFNKI<br>27 TPNLAEFAFS LYRQLAHQS**N** STNIFFSPVS IATAFAMLSL GTKADTHDEI<br>77 LEGLNF**N**LTE IPEAQIHEGF QELLRTLNQP DSQLQLTTGN GLFLSEGLKL<br>127 VDKFLEDVKK LYHSEAFTVN FGDTEEAKKQ INDYVEKGTQ GKIVDLVKEL<br>177 DRDTVFALVN YIFFKGKWER PFEVKDTEEE DFHVDQVTTV KVPMMKRLGM<br>227 FNIQHCKKLS SWVLLMKYLG **N**ATAIFFLPD EGKLQHLENE LTHDIITKFL<br>277 ENEDRRSASL HLPKLSITGT YDLKSVLGQL GITKVFSNGA DLRGHRGGTP<br>327 EALQGRA |

| Construct | Residue | Sequence |
|---|---|---|
| **N-terminally tagged *Null* Hong Kong (N-NHK)**[2] | | 123 |
| | -24 | MPSSVSWGIL LLAGLCCLVP VSLAEDPEQK LISEEDLNSA VDHHHHHHEN |
| | | 4 |
| | | LYFQGQGDAA QKTDTSHHDQ DHPTFNKITP NLAEFAFSLY RQLAHQSNST |
| | 49 | NIFFSPVSIA TAFAMLSLGT KADTHDEILE GLNFNLTEIP EAQIHEGFQE |
| | 99 | LLRTLNQPDS QLQLTTGNGL FLSEGLKLVD KFLEDVKKLY HSEAFTVNFG |
| | 149 | DTEEAKKQIN DYVEKGTQGK IVDLVKELDR DTVFALVNYI FFKGKWERPF |
| | 199 | EVKDTEEEDF HVDQVTTVKV PMMKRLGMFN IQHCKKLSSW VLLMKYLGNA |
| | 249 | TAIFFLPDEG KLQHLENELT HDIITKFLEN EDRRSASLHL PKLSITGTYD |
| | 299 | LKSVLGQLGI TKVFSNGADL RGHRGGTPEA LQGRA |

| Construct | Residue | Sequence |
|---|---|---|
| **Full-length AAT** | | 123 |
| | -24 | MPSSVSWGIL LLAGLCCLVP VSLAEDPEQK LISEEDLNSA VDHHHHHHEN |
| | | 4 |
| | | LYFQGQGDAA QKTDTSHHDQ DHPTFNKITP NLAEFAFSLY RQLAHQSNST |
| | 49 | NIFFSPVSIA TAFAMLSLGT KADTHDEILE GLNFNLTEIP EAQIHEGFQE |
| | 99 | LLRTLNQPDS QLQLTTGNGL FLSEGLKLVD KFLEDVKKLY HSEAFTVNFG |
| | 149 | DTEEAKKQIN DYVEKGTQGK IVDLVKELDR DTVFALVNYI FFKGKWERPF |
| | 199 | EVKDTEEEDF HVDQVTTVKV PMMKRLGMFN IQHCKKLSSW VLLMKYLGNA |
| | 249 | TAIFFLPDEG KLQHLENELT HDIITKFLEN EDRRSASLHL PKLSITGTYD |
| | 299 | LKSVLGQLGI TKVFSNGADL SGVTEEAPLK LSKAVHKAVL TIDEKGTEAA |
| | 349 | GAMFLEAIPM SIPPEVKFNK PFVFLMIEQN TKSPLFMGKV VNPTQK |

[1]In N-terminally tagged constructs, to ensure proper cleavage of the signal sequence, the first three amino acids of the mature AAT sequence precede the N-terminal tag.

[2] Note that the originally reported NHK sequence was in the AAT M2 background which has an Arg101His mutation relative to AAT M1V (2). All versions of NHK used in these studies are in the M1V background (Arg101).

**Table S3.** Characteristics of the $R_g$ distribution for each HDXer cluster.

| Cluster | Number of Structures | Mean $R_g$ (Å) | Standard Deviation (Å) | Skewness |
|---|---|---|---|---|
| 1 | 2932 | 17.6 | 0.5 | 0.59 |
| 2 | 1267 | 16.8 | 0.1 | 0.03 |
| 3 | 2856 | 17.5 | 0.6 | 1.21 |
| 4 | 1299 | 16.7 | 0.1 | -0.06 |
| 5 | 1886 | 17.2 | 0.1 | 0.13 |
| 6 | 695 | 18.2 | 0.7 | 1.19 |

**Table S4.** DNA primer sequences with mutations in bold.

| Purpose | Direction | DNA Primer Sequence |
|---|---|---|
| | | **AAT fragment expression in *Escherichia coli*** |
| Site Directed Mutagenesis (SDM): Cys232Ser | Forward | 5′ GTTTAACATCCAGCAC**TCT**AAGAAGCTGTCCAGC 3′ |
| | Reverse | 5′ GCTGGACAGCTTCTT**AGA**GTGCTGGATGTTAAAC 3′ |
| AAT190: introduce a stop codon at 191 | Forward | 5′ GAATTACATCTTCTTT**T**AAGGCAAATGGGAGAC 3′ |
| | Reverse | 5′ GTCTCCCATTTGCCTTAAA**A**GAAGATGTAATTC 3′ |
| AAT290: introduce a stop codon at 291 | Forward | 5′ GCTTACATTTACCCAAA**TAA**TCCATTACTGGAACC 3′ |
| | Reverse | 5′ GGTTCCAGTAATGGA**TTA**TTTGGGTAAATGTAAGC 3′ |
| AAT323: introduce a stop codon at 324 | Forward | 5′ CGGGGTCACAGAG**TAA**GCACCCCTG 3′ |
| | Reverse | 5′ CAGGGGTGC**TTA**CTCTGTGACCCCG 3′ |
| | | **Full-length AAT expression in *Escherichia coli:* pEAT8-137 plasmid change AAT M2 sequence to AAT M1V sequence** |
| SDM His101Arg | Forward | 5′ CCAGGAACTCCTCC**G**TACCCTCAACCAGC 3′ |
| | Reverse | 5′ GCTGGTTGAGGGTA**C**GGAGGAGTTCCTGG 3′ |
| SDM Asp376Glu | Forward | 5′ CCTTTGTCTTCTTAATGATTGA**A**CAAAATACCAAGTCTCC 3′ |
| | Reverse | 5′ GGAGACTTGGTATTTTG**T**TCAATCATTAAGAAGACAAAGG 3′ |
| | | **Insert the N-terminal tag for AAT expression in mammalian cells** |
| Insert the N-terminal tags | Forward[1] | 5′ GTCTCCCTGGCTGAGGATCCCGAACAAAAACTCATCTCAGAAGAG 3′ |
| | Reverse[2] | 5′ GGCAGCATCTCCCTGCCCCTGGAAGTAGAGATTCTCATGATGATGATG 3′ |
| Amplify the pcDNA vector backbone including AAT M1V | Forward | 5′ CATCATCATCATGAGAATCTCTACTTCCAGGGGCAGGGAGATGCTGCC 3′ |
| | Reverse | 5′ CTCTTCTGAGATGAGTTTTTGTTCGGATCCTCAGCCAGGGAGAC 3′ |

[1]The signal sequence, which is complementary to the pcDNA plasmid containing AAT M1V, is in bluish green, the first three codons of mature AAT are in black and the codons for the Myc tag are in vermillion.

[2]Codon 4 to 8 of mature AAT M1V in black, codons for the TEV cleavage site in light blue and the codons for the 6XHis tag in vermillion.

**Table S5.** Protein sequences for full length AAT M1V expressed in *E. coli* as a soluble protein using a modified pEAT8-137 plasmid (3). The serine residue mutated to phenylalanine in the Siiyama variant (S53F), is highlighted in yellow and the single cysteine, 232, which is mutated to serine is in purple.

| AAT Construct | Amino Acid Sequence* | Molecular Weight (kDa) | $\varepsilon_{280\ nm}$ ($M^{-1}\ cm^{-1}$) |
|---|---|---|---|
| Full-length AAT C232S | 1 MDPQGDAAQK TDTSHHDQDH PTFNKITPNL<br>31 AEFAFSLYRQ LAHQSNSTNI FF**S**PVSIATA<br>61 FAMLSLGTKA DTHDEILEGL NFNLTEIPEA<br>91 QIHEGFQELL RTLNQPDSQL QLTTGNGLFL<br>121 SEGLKLVDKF LEDVKKLYHS EAFTVNFGDT<br>151 EEAKKQINDY VEKGTQGKIV DLVKELDRDT<br>181 VFALVNYIFF KGKWERPFEV KDTEEEDFHV<br>211 DQVTTVKVPM MKRLGMFNIQ H**S**KKLSSWVL<br>241 LMKYLGNATA IFFLPDEGKL QHLENELTHD<br>271 IITKFLENED RRSASLHLPK LSITGTYDLK<br>301 SVLGQLGITK VFSNGADLSG VTEEAPLKLS<br>331 KAVHKAVLTI DEKGTEAAGA MFLEAIPMSI<br>361 PPEVKFNKPF VFLMIEQNTK SPLFMGKVVN<br>391 PTQK | 44,310 | 23,484** |

*In this full-length construct, the N-terminal glutamic acid residue is mutated to methionine (E1M) (4).

** $E_{280\ nm}^{0.1\%} = 0.53$ (5)

**Table S6.** Hydrogen-deuterium exchange mass spectrometry (HDX-MS) data quality (6) for AAT190 and full-length AAT.

| Data set | AAT190 Biological Replicate 1 | AAT190 Biological Replicate 2 | Full Length AAT |
|---|---|---|---|
| **HDX reaction conditions** | 10µl of 8µM AAT190 diluted in 90µl of $^2H_2O$ Buffer (50 mM potassium phosphate, 150 mM NaCl, pD 7.4) | 10µl of 8µM AAT190 diluted in 90µl of $^2H_2O$ Buffer (50 mM potassium phosphate, 150 mM NaCl, pD 7.4) | 10µl of 8µM full-length AAT diluted in 90µl of $^2H_2O$ Buffer (50 mM potassium phosphate, 150 mM NaCl, pD 7.4) |
| **HDX incubation times** | 10 sec, 1min, 10min, 1hr, 2hr | 10 sec, 1min, 10min, 1hr, 2hr | 10 sec, 1min, 10min, 1hr, 2hr |
| **HDX controls** | Un-deuterated control, fully deuterated control | Un-deuterated control, fully deuterated control | Un-deuterated control, fully deuterated control |
| **Back exchange (based on fully deuterated controls)** | 31 +/- 9% | 31 +/- 9% | 35 +/- 7% |
| **Number of peptides** | 29 | 29 | 66 |
| **Sequence coverage** | 96.3% | 96.3% | 91.6% |
| **Average peptide length/redundancy** | 12.7 /2.62 | 12.7/2.62 | 13.1/2.36 |
| **Replicates (biological or technical)** | Three technical | Three technical | Three technical |
| **Repeatability** | 1.2% standard deviation across technical replicates | 1.6% standard deviation across technical replicates | 2.2% standard deviation across technical replicates |

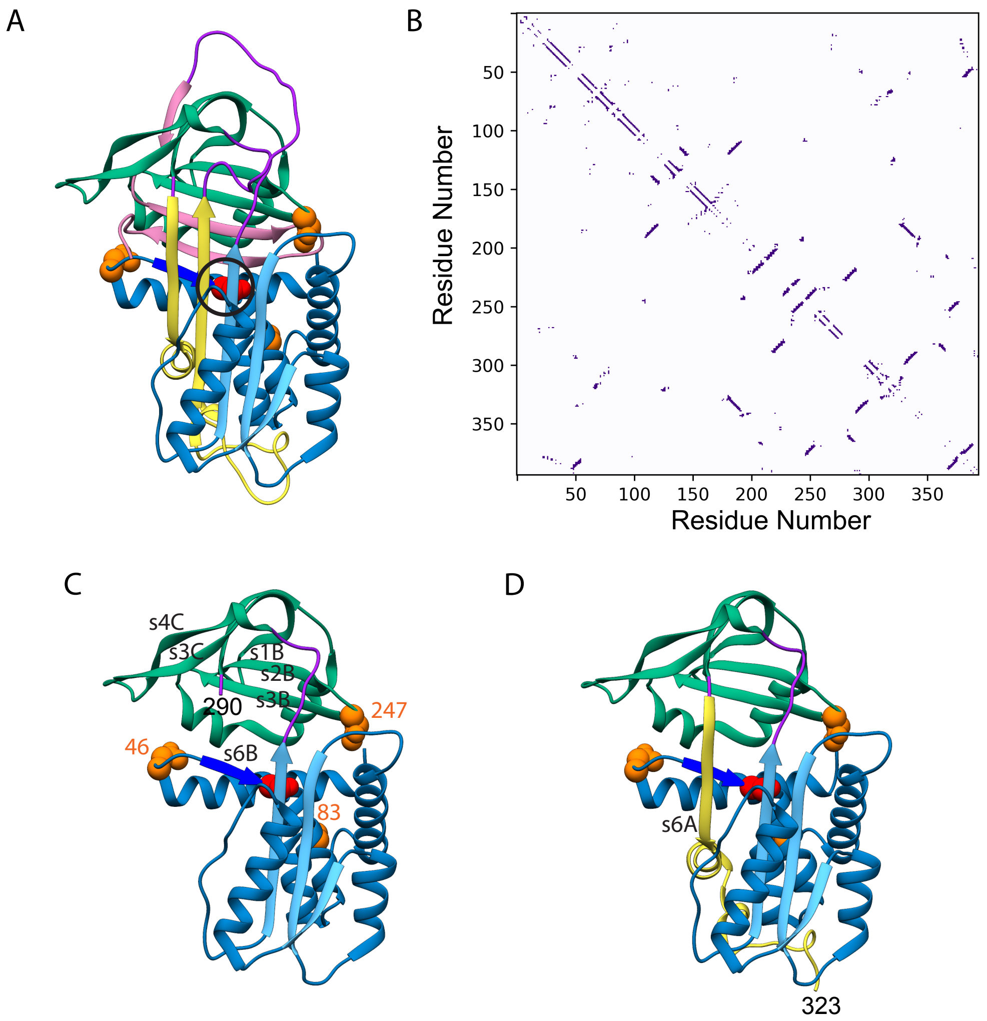


**Figure S1.** Full-length AAT and the structures of the 1-290 (AAT290) and 1-323 (AAT323) fragments in the context of the full-length protein. (A) As also shown in Figure 1A, the x-ray crystal structure of full-length AAT (PDB:1QLP (7)).The N- and C-terminal regions of the αβ domain are in blue and yellow, respectively; the N-terminal regions of the mainly β domain are in bright blue (strand 6B) and green with the C-terminal region in pink; loops connecting the two domains, including the long functionally required RCL, are in purple; and asparagines that are glycosylated in the ER are in orange spacefill. The black circle indicates the functionally relevant shutter region containing the Siiyama (Ser53Phe) mutation (red spacefill). (B) Full-length AAT contact map from PDB:1QLP constructed using a cut-off of 7.5 Å for alpha carbons. The contact map includes the disordered N-terminus (residues 1-22) which was added using the CHARMM-GUI (8). (C) Structure of AAT290 in the context of the full-length serpin. (D) Structure of AAT323 in the context of the full-length serpin.

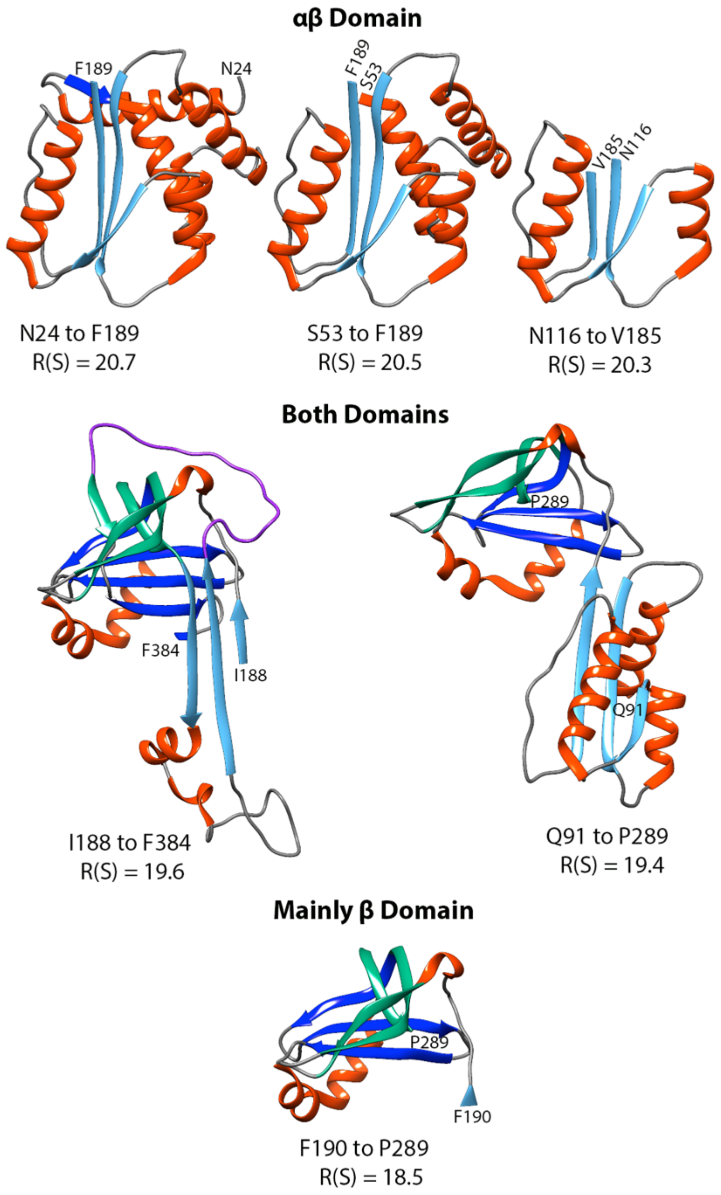


**Figure S2.** RAFT prediction (9) of sequential autonomous folding units (AFUs). The RAFT predictions showing the structures, in the context of full-length $\alpha_1$-antitrypsin (AAT) and the RAFT scores, R(S). The helices are in orange; β sheet A, which defines the αβ domain, is in sky blue; β sheets B and C which define the mainly β domain, are in dark blue and bluish green, respectively and the RCL is in purple. The calculations are courtesy of Dr. Kael F. Fischer.

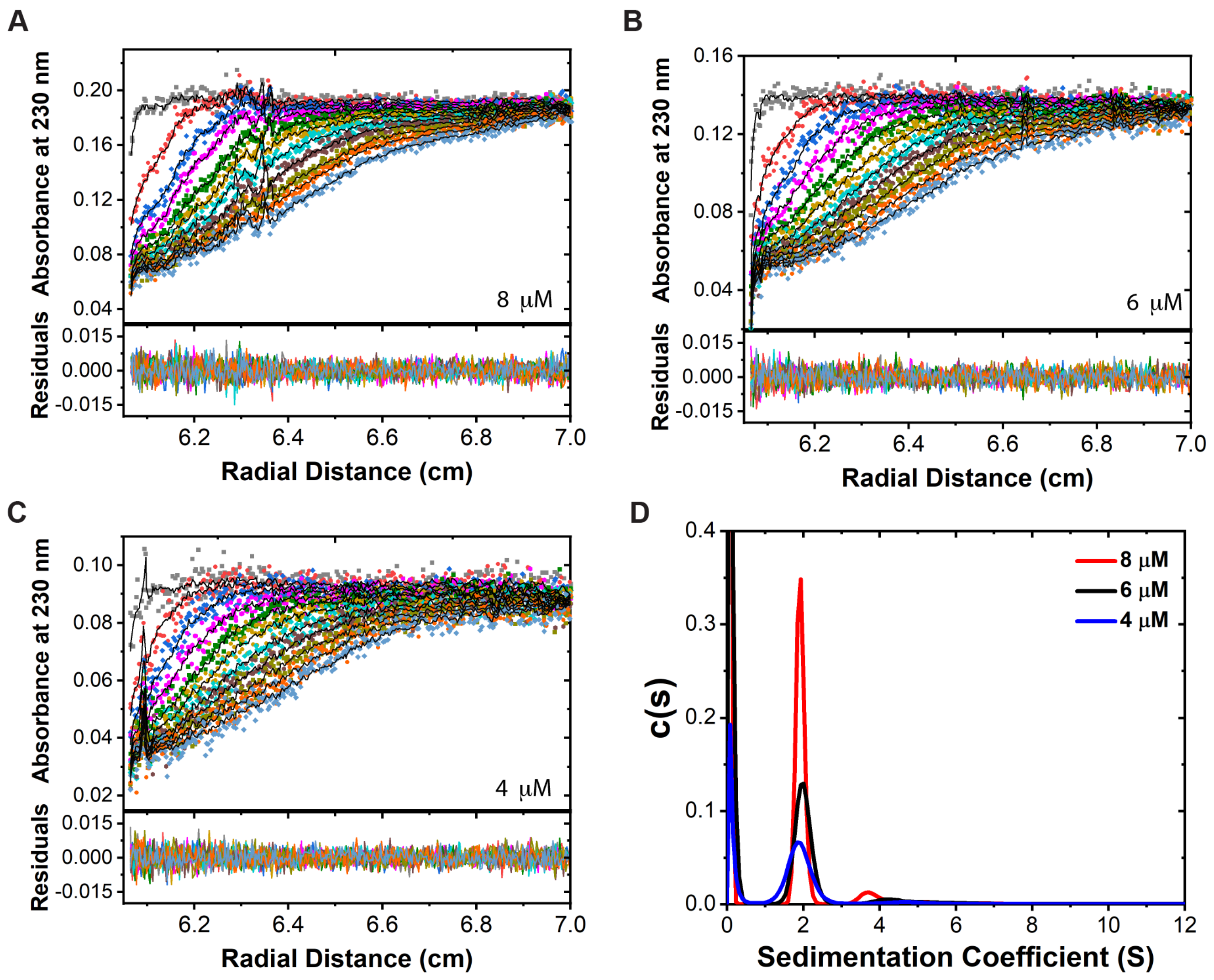


**Figure S3.** AAT190 is mainly monomeric. Sedimentation velocity data (colored points), fits (black lines) and residuals for AAT190 (A) 8 µM, (B) 6 µM and (C) 4 µM in 50 mM potassium phosphate. pH 7.4. (D) Distributions of sedimentation coefficients for each concentration. The peak near 0 arises from a population of small molecules much smaller than the AAT190 fragment. Data analysis was performed using SEDFIT (10, 11).

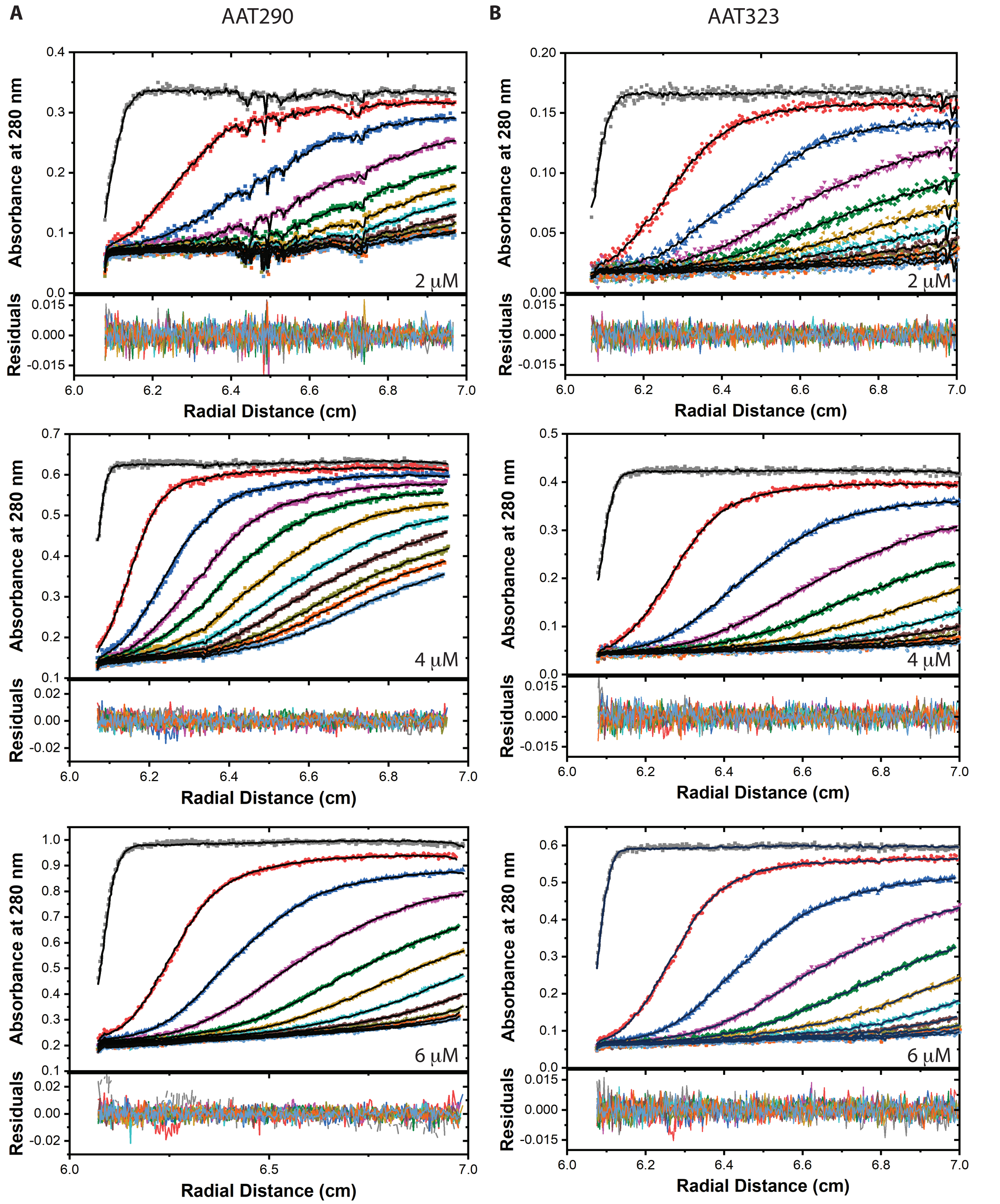


**Figure S4.** AUC sedimentation velocity data, fits and residuals for (A) AAT290 and (B) AAT323 for 2 μM (top), 4 μM (middle) and 6 μM (bottom). Every tenth scan is plotted with the data as colored squares and the fits as black lines. For the sedimentation coefficient distributions see Figure 2 in the main text. Data analysis was performed using SEDFIT (10, 11).

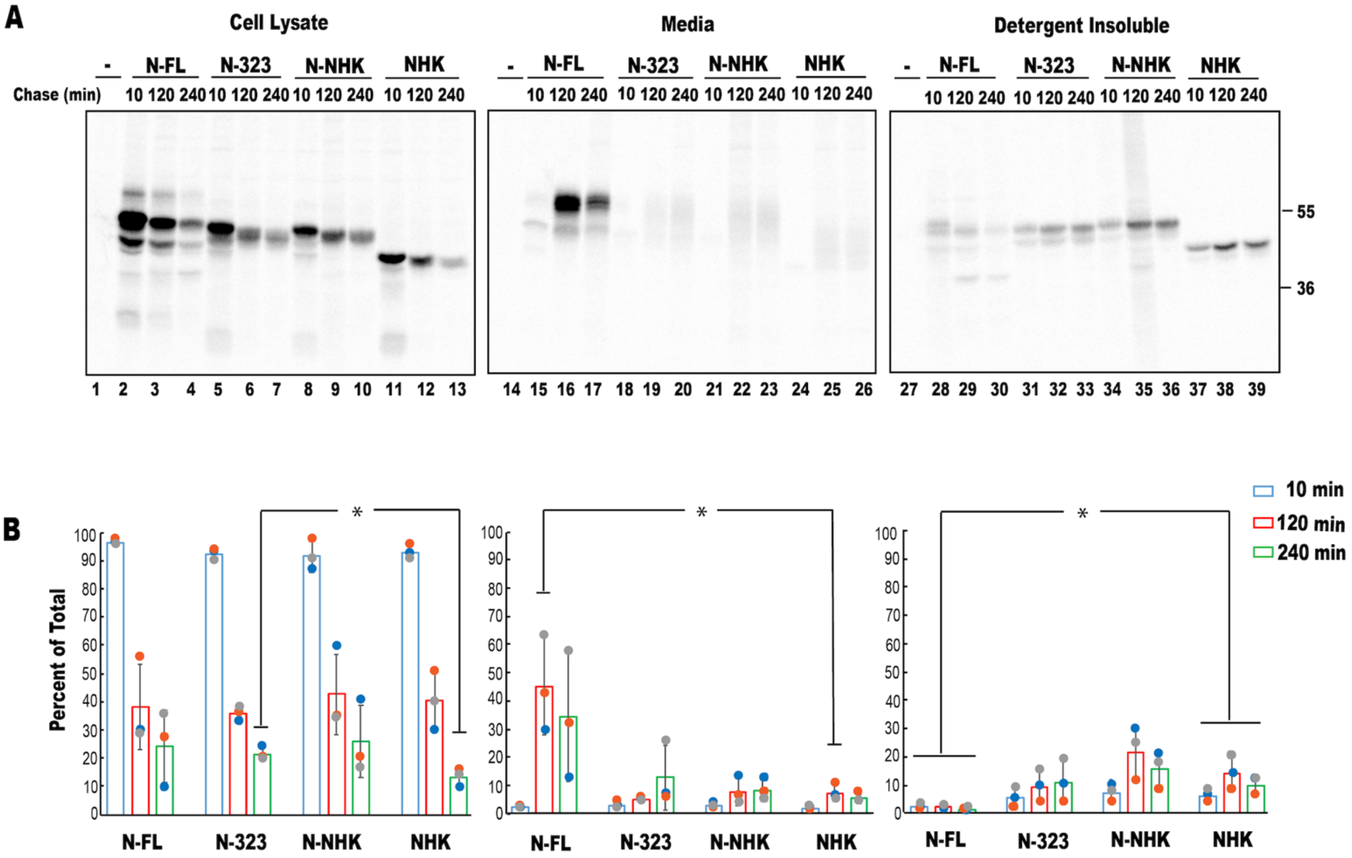


**Figure S5.** AAT323 behaves similarly to the AAT mutant *null* Hong Kong (NHK) and is targeted for degradation. (A) CHO cells were grown for 20 hr, radiolabeled using a 30 min pulse and chased for 10, 120 and 240 min using regular growth media. NHK with no tag, the variant often used as an ERAD substrate serves as a control. All other constructs, full-length AAT (N-FL), AAT323 (N-323) and N-NHK were epitope tagged at the N-terminus. For all variants, the amount of protein in the cell lysate decreased over time (lanes 2-13). For full-length AAT the decrease in the amount of protein in the cell lysate correlates with secretion into the media (lanes 15-17) with little accumulation in the detergent insoluble fraction (lanes 29-30). While some secretion and accumulation in the detergent insoluble fraction is observed over time for NHK (lanes 24-26 and 37-39, respectively), N-NHK (lanes 21-23 and 34-36, respectively) and AAT323 (N-323, lanes 18-20 and 31-33, respectively), these small increases do not account for the large decrease in protein abundance observed in the cell lysate. Results from transfection with the empty plasmid (-) are shown in lanes 1, 14 and 27. (B) Quantification of the results from three biological replicates (blue, orange and gray circles). For each AAT variant, the data are normalized relative to the total amount of protein (cell lysate plus media plus detergent-insoluble) at the 10 min chase time point. The mean percent of protein is indicated by the bars with chase time points at 10 min, 120 min and 240 min shown in blue, red and green respectively. The error bars show the standard deviation. Unpaired t-tests were performed between C-terminally tagged NHK and each variant for every condition and chase time, asterisks indicate $p<0.05$.

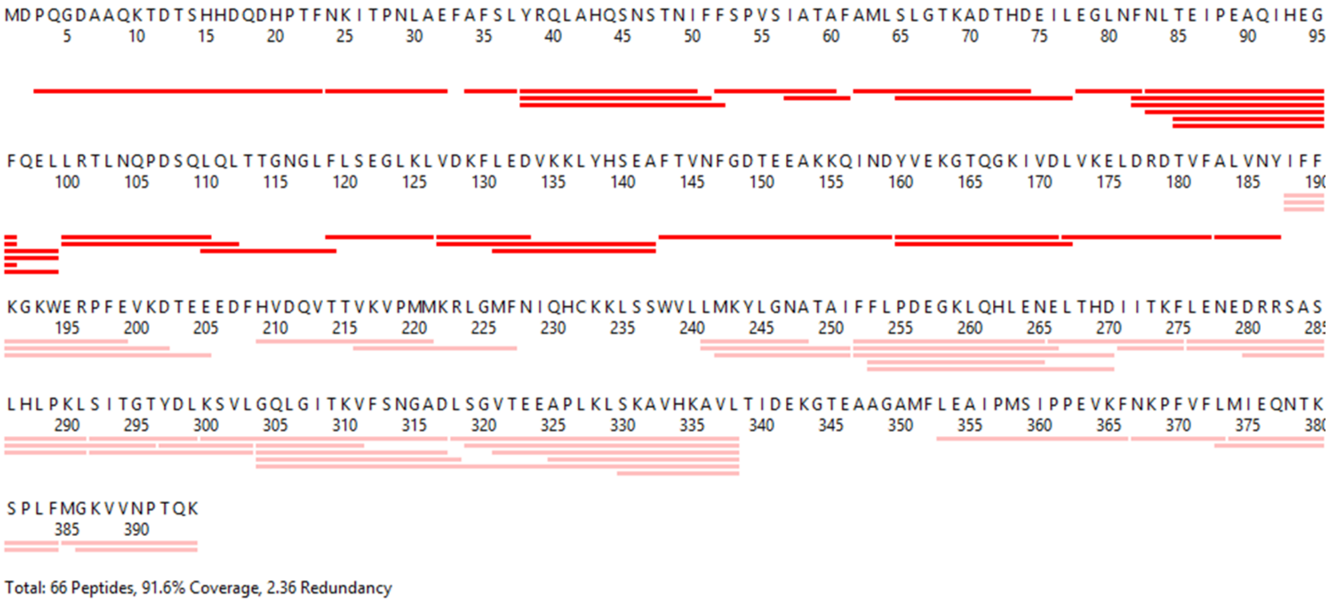


**Figure S6**. Hydrogen-deuterium exchange mass spectrometry (HDX-MS) sequence coverage of full-length AAT. Horizontal bars represent individual peptides that were identified by the ProteinLynx Global server for which both deuterium uptake data and a fully deuterated control were successfully acquired. Peptides identified in the full length and AAT190 context are shown in red, peptides identified only in the full-length context are shown in pink. For full-length AAT there are 66 peptides total resulting in 91.6% coverage and a redundancy of 2.36.

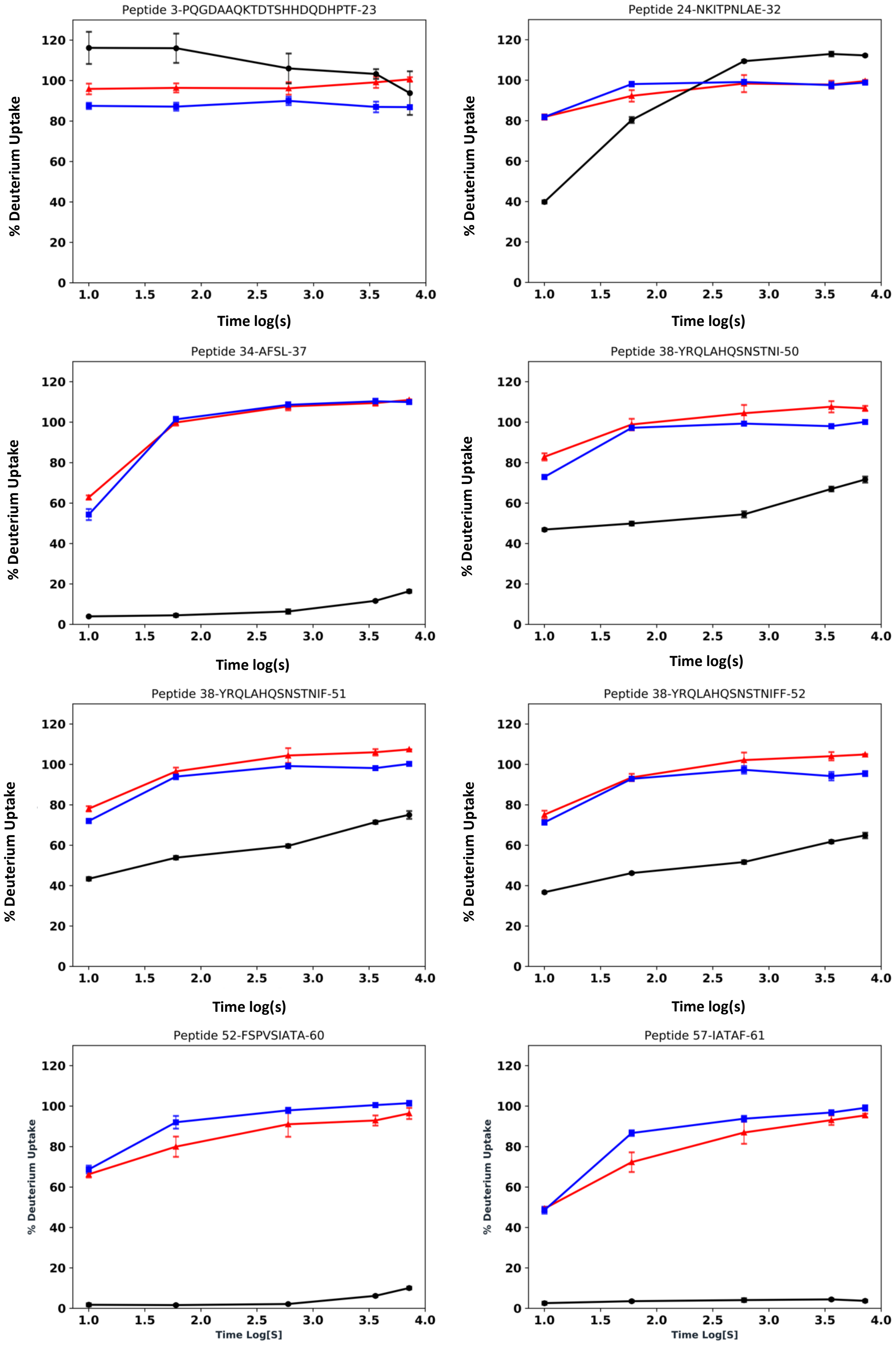
Peptide 3-PQGDAAQKTDTSHHDQDHPTF-23
Peptide 24-NKITPNLAE-32
Peptide 34-AFSL-37
Peptide 38-YRQLAHQSNSTNI-50
Peptide 38-YRQLAHQSNSTNIF-51
Peptide 38-YRQLAHQSNSTNIFF-52
Peptide 52-FSPVSIATA-60
Peptide 57-IATAF-61
% Deuterium Uptake
Time log(s)
Time Log[S]

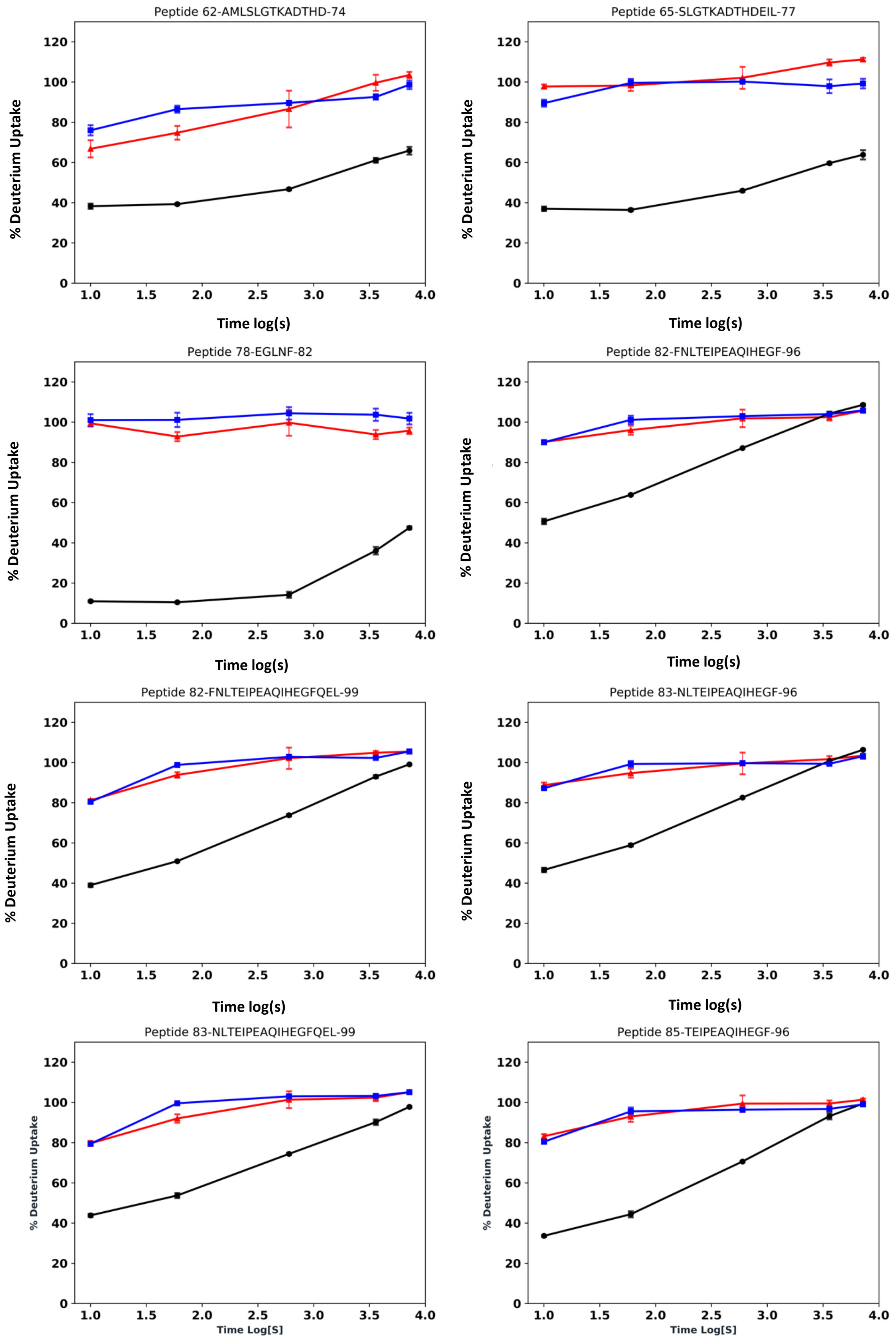
Peptide 62-AMLSLGTKADTHD-74
Peptide 65-SLGTKADTHDEIL-77
Peptide 78-EGLNF-82
Peptide 82-FNLTEIPEAQIHEGF-96
Peptide 82-FNLTEIPEAQIHEGFQEL-99
Peptide 83-NLTEIPEAQIHEGF-96
Peptide 83-NLTEIPEAQIHEGFQEL-99
Peptide 85-TEIPEAQIHEGF-96
% Deuterium Uptake
Time log(s)
Time Log[S]

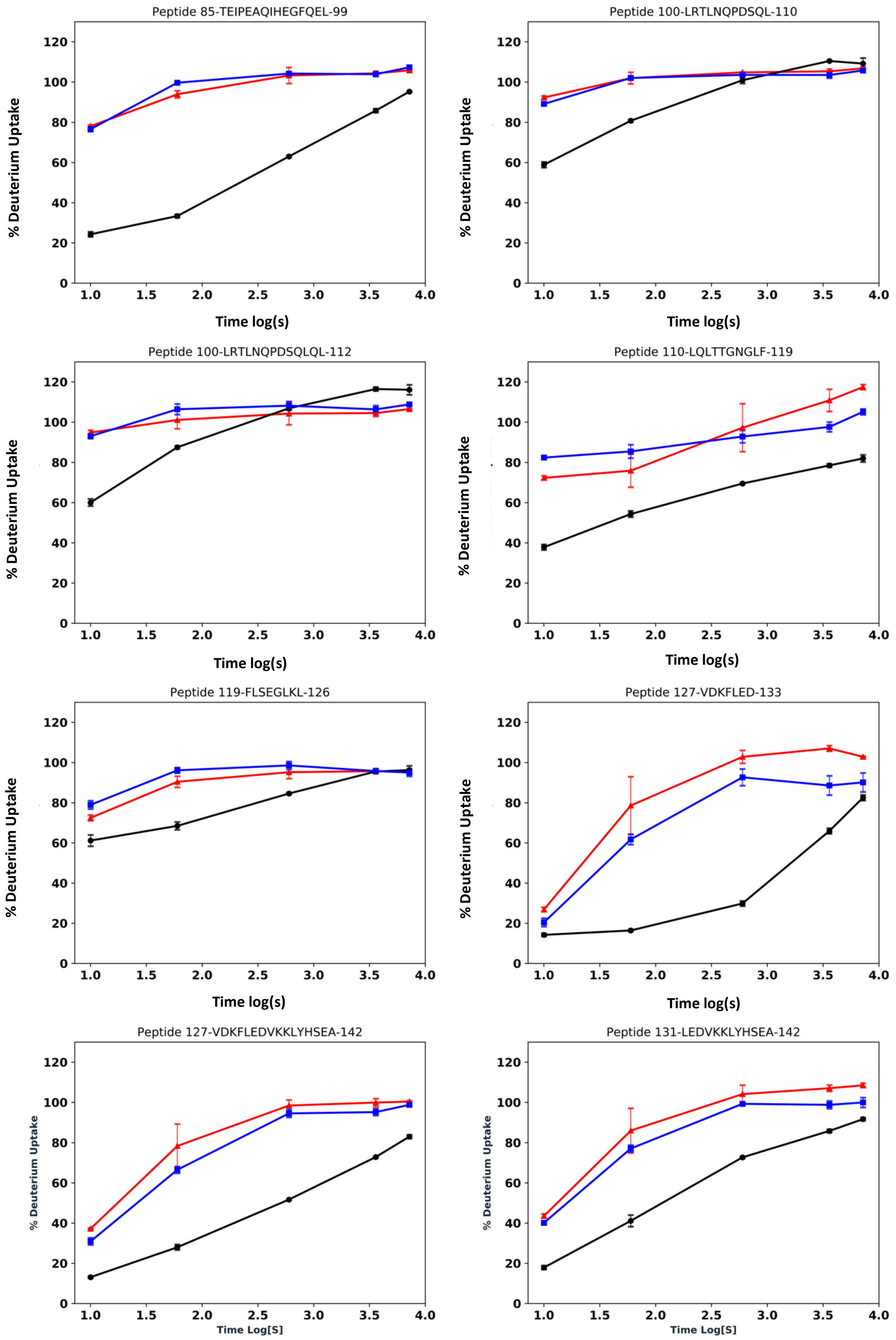
Peptide 85-TEIPEAQIHEGFQEL-99
Peptide 100-LRTLNQPDSQL-110
Peptide 100-LRTLNQPDSQLQL-112
Peptide 110-LQLTTGNGLF-119
Peptide 119-FLSEGLKL-126
Peptide 127-VDKFLED-133
Peptide 127-VDKFLEDVKKLYHSEA-142
Peptide 131-LEDVKKLYHSEA-142
% Deuterium Uptake
Time log(s)
Time Log[S]

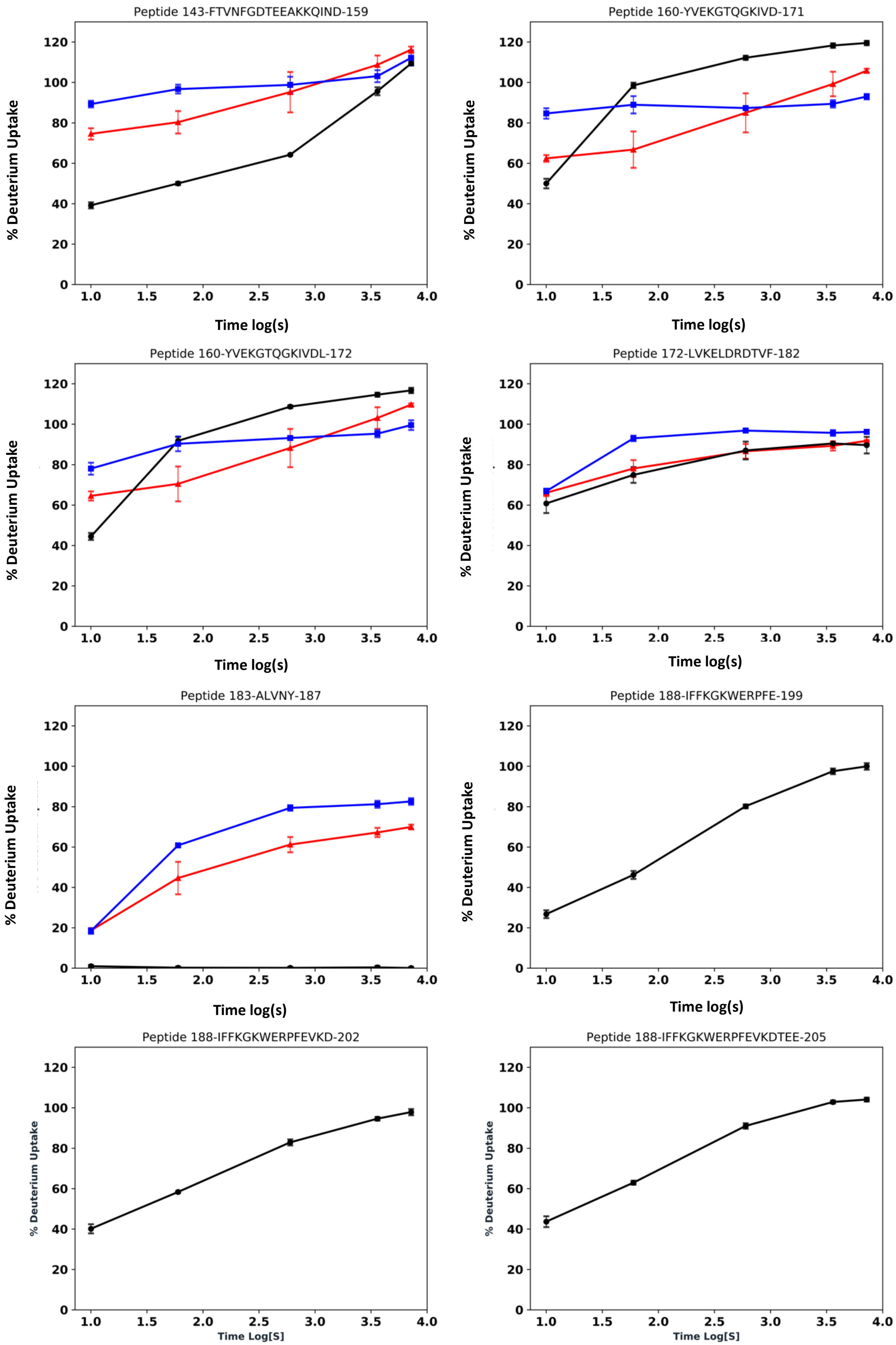
Peptide 143-FTVNFGDTEEAKKQIND-159
% Deuterium Uptake
Time log(s)
Peptide 160-YVEKGTQGKIVD-171
% Deuterium Uptake
Time log(s)
Peptide 160-YVEKGTQGKIVDL-172
% Deuterium Uptake
Time log(s)
Peptide 172-LVKELDRDTVF-182
% Deuterium Uptake
Time log(s)
Peptide 183-ALVNY-187
% Deuterium Uptake
Time log(s)
Peptide 188-IFFKGKWERPFE-199
% Deuterium Uptake
Time log(s)
Peptide 188-IFFKGKWERPFEVKD-202
% Deuterium Uptake
Time Log[S]
Peptide 188-IFFKGKWERPFEVKDTEE-205
% Deuterium Uptake
Time Log[S]

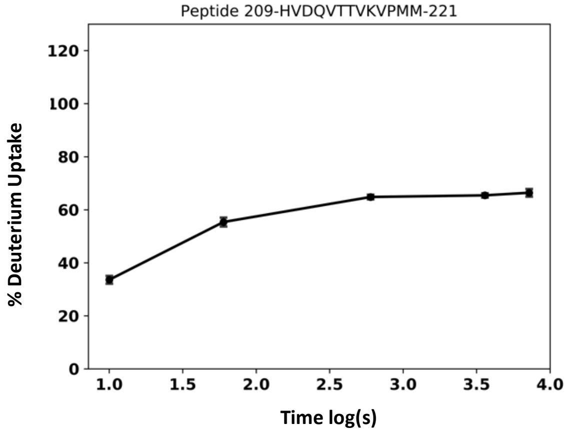
Peptide 209-HVDQVTTVKVPMM-221
% Deuterium Uptake
0 20 40 60 80 100 120
1.0 1.5 2.0 2.5 3.0 3.5 4.0
Time log(s)

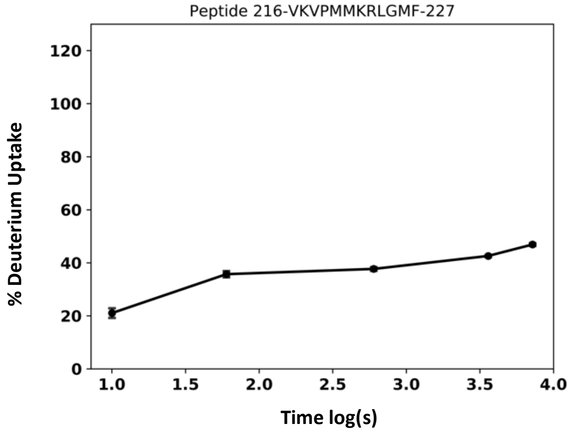
Peptide 216-VKVPMMKRLGMF-227
% Deuterium Uptake
0 20 40 60 80 100 120
1.0 1.5 2.0 2.5 3.0 3.5 4.0
Time log(s)

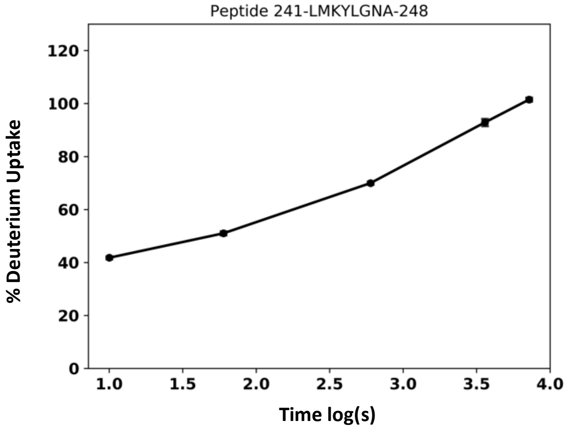
Peptide 241-LMKYLGNA-248
% Deuterium Uptake
0 20 40 60 80 100 120
1.0 1.5 2.0 2.5 3.0 3.5 4.0
Time log(s)

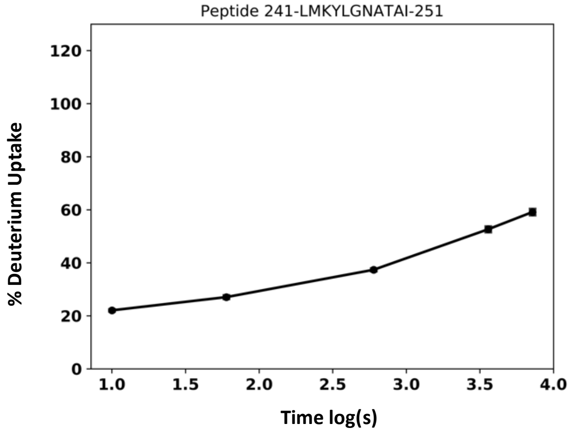
Peptide 241-LMKYLGNATAI-251
% Deuterium Uptake
0 20 40 60 80 100 120
1.0 1.5 2.0 2.5 3.0 3.5 4.0
Time log(s)

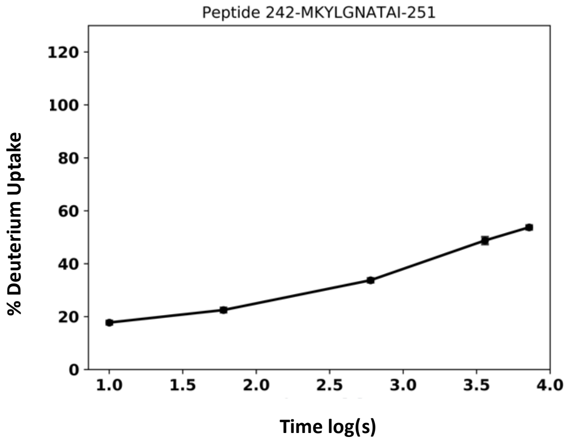
Peptide 242-MKYLGNATAI-251
% Deuterium Uptake
0 20 40 60 80 100 120
1.0 1.5 2.0 2.5 3.0 3.5 4.0
Time log(s)

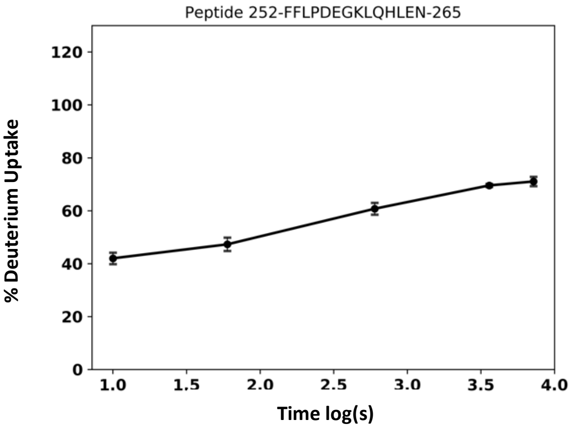
Peptide 252-FFLPDEGKLQHLEN-265
% Deuterium Uptake
0 20 40 60 80 100 120
1.0 1.5 2.0 2.5 3.0 3.5 4.0
Time log(s)

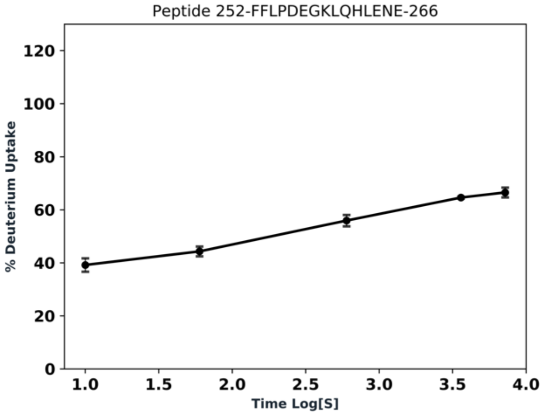
Peptide 252-FFLPDEGKLQHLENE-266
% Deuterium Uptake
0 20 40 60 80 100 120
1.0 1.5 2.0 2.5 3.0 3.5 4.0
Time Log[S]

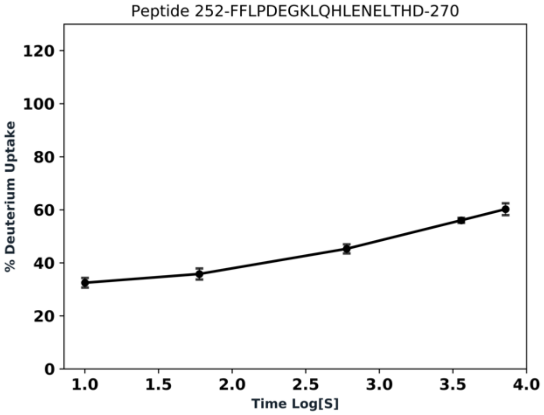
Peptide 252-FFLPDEGKLQHLENELTHD-270
% Deuterium Uptake
0 20 40 60 80 100 120
1.0 1.5 2.0 2.5 3.0 3.5 4.0
Time Log[S]

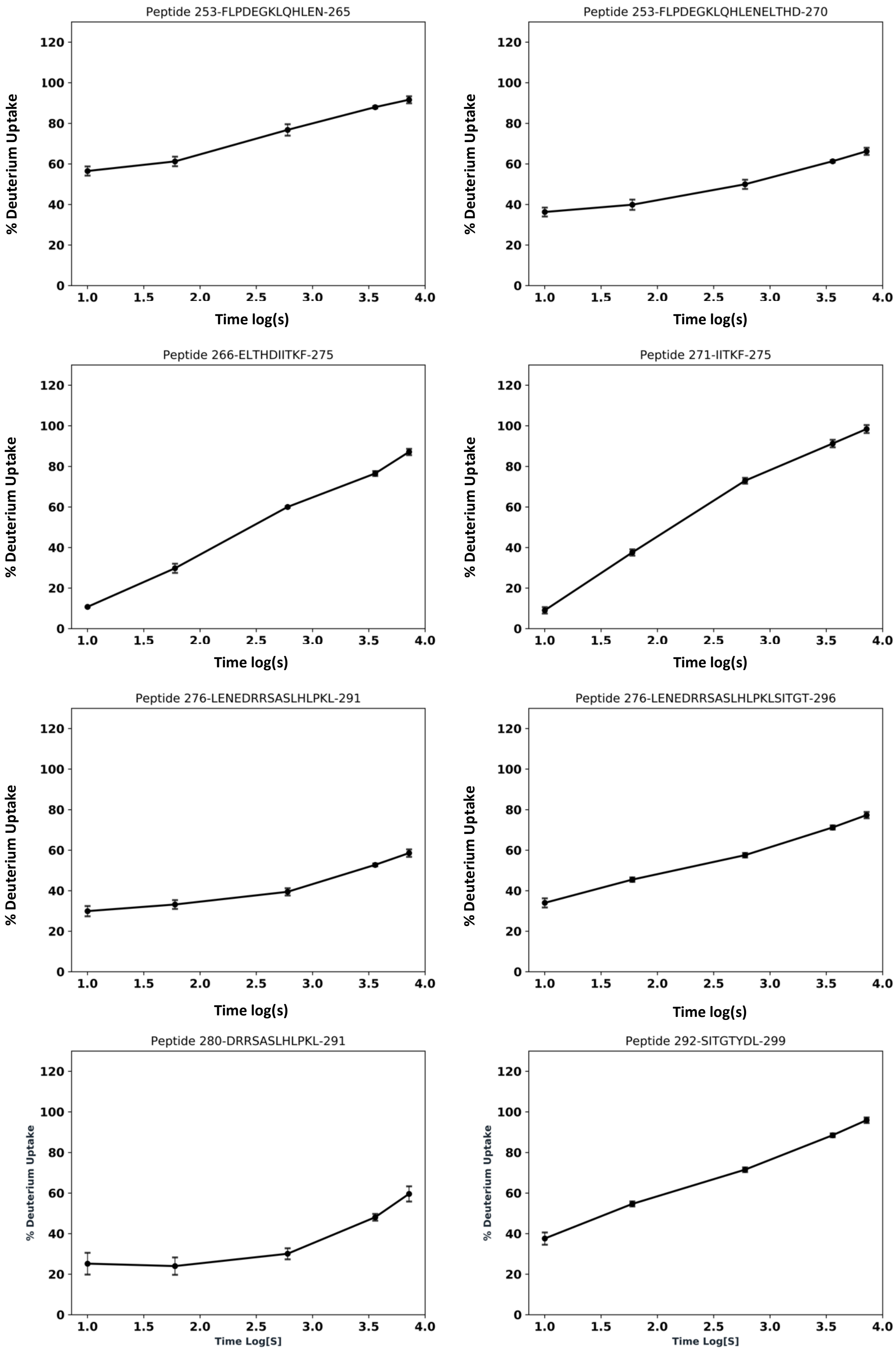
Peptide 253-FLPDEGKLQHLEN-265
Peptide 253-FLPDEGKLQHLENELTHD-270
Peptide 266-ELTHDIITKF-275
Peptide 271-IITKF-275
Peptide 276-LENEDRRSASLHLPKL-291
Peptide 276-LENEDRRSASLHLPKLSITGT-296
Peptide 280-DRRSASLHLPKL-291
Peptide 292-SITGTYDL-299
% Deuterium Uptake
Time log(s)
Time Log[S]
0
20
40
60
80
100
120
1.0
1.5
2.0
2.5
3.0
3.5
4.0

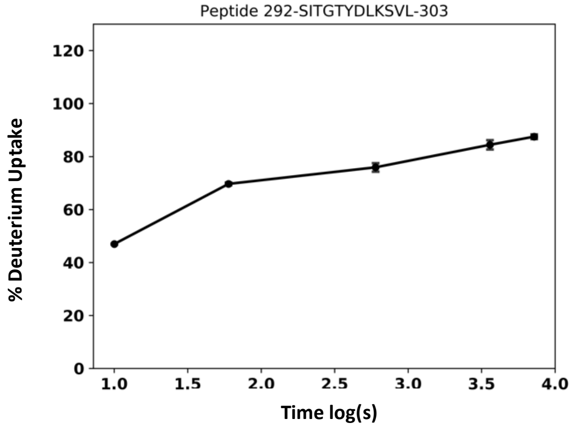
Peptide 292-SITGTYDLKSVL-303
% Deuterium Uptake
Time log(s)

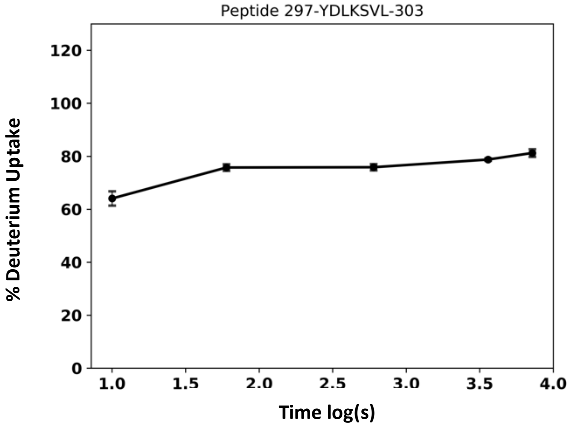
Peptide 297-YDLKSVL-303
% Deuterium Uptake
Time log(s)

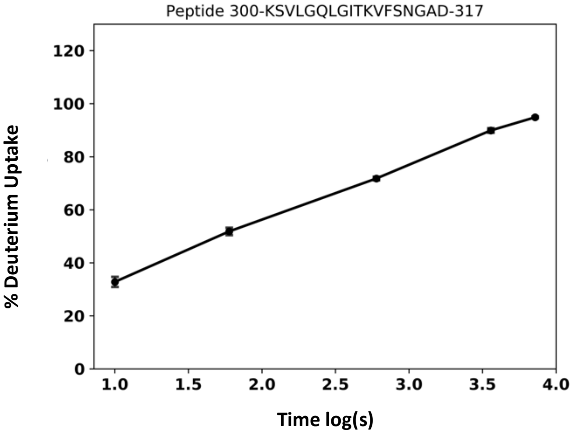
Peptide 300-KSVLGQLGITKVFSNGAD-317
% Deuterium Uptake
Time log(s)

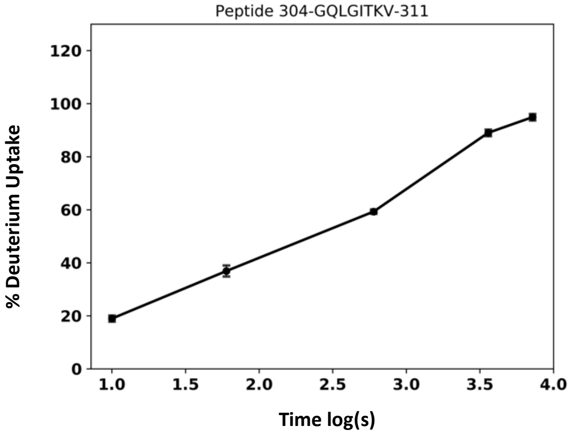
Peptide 304-GQLGITKV-311
% Deuterium Uptake
Time log(s)

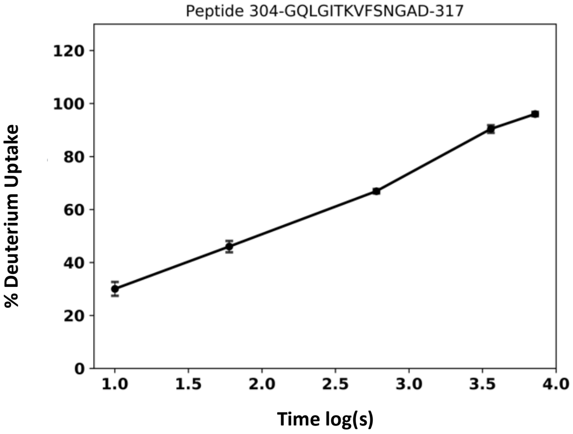
Peptide 304-GQLGITKVFSNGAD-317
% Deuterium Uptake
Time log(s)

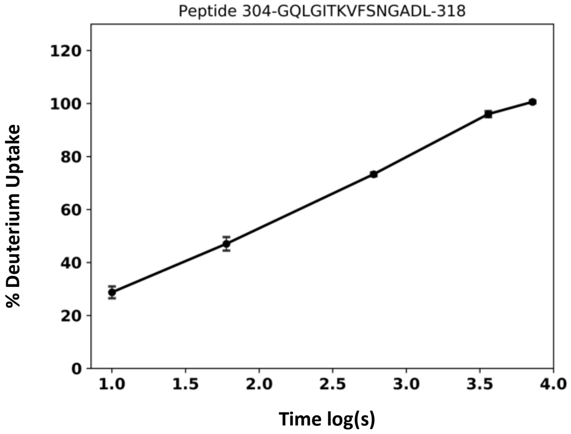
Peptide 304-GQLGITKVFSNGADL-318
% Deuterium Uptake
Time log(s)

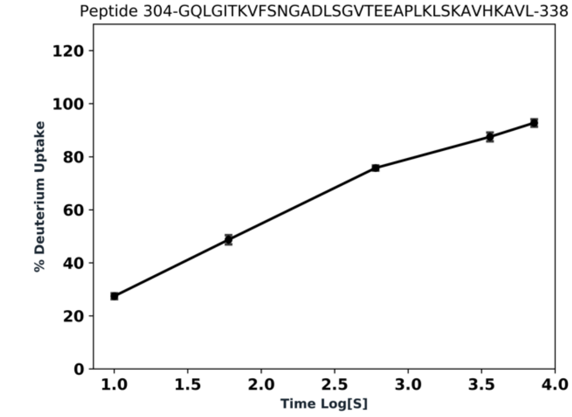
Peptide 304-GQLGITKVFSNGADLSGVTEEAPLKLSKAVHKAVL-338
% Deuterium Uptake
Time Log[S]

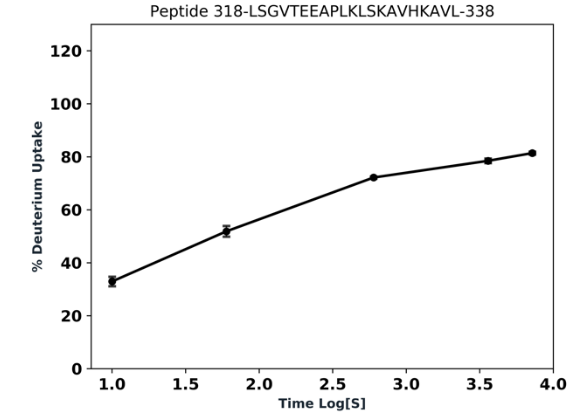
Peptide 318-LSGVTEEAPLKLSKAVHKAVL-338
% Deuterium Uptake
Time Log[S]

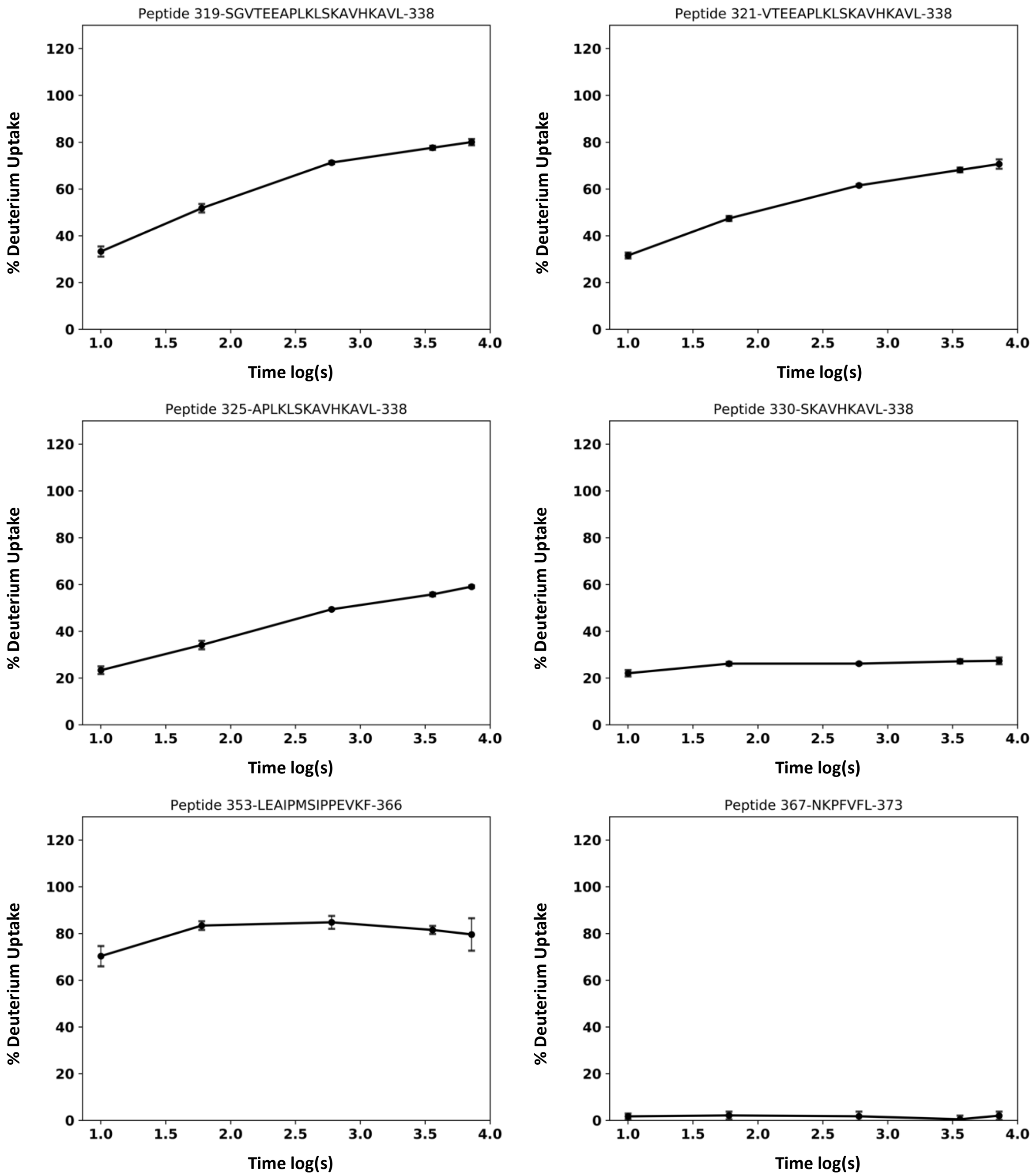
Peptide 319-SGVTEEAPLKLSKAVHKAVL-338
Peptide 321-VTEEAPLKLSKAVHKAVL-338
Peptide 325-APLKLSKAVHKAVL-338
Peptide 330-SKAVHKAVL-338
Peptide 353-LEAIPMSIPPEVKF-366
Peptide 367-NKPFVFL-373
% Deuterium Uptake
Time log(s)
0
20
40
60
80
100
120
1.0
1.5
2.0
2.5
3.0
3.5
4.0

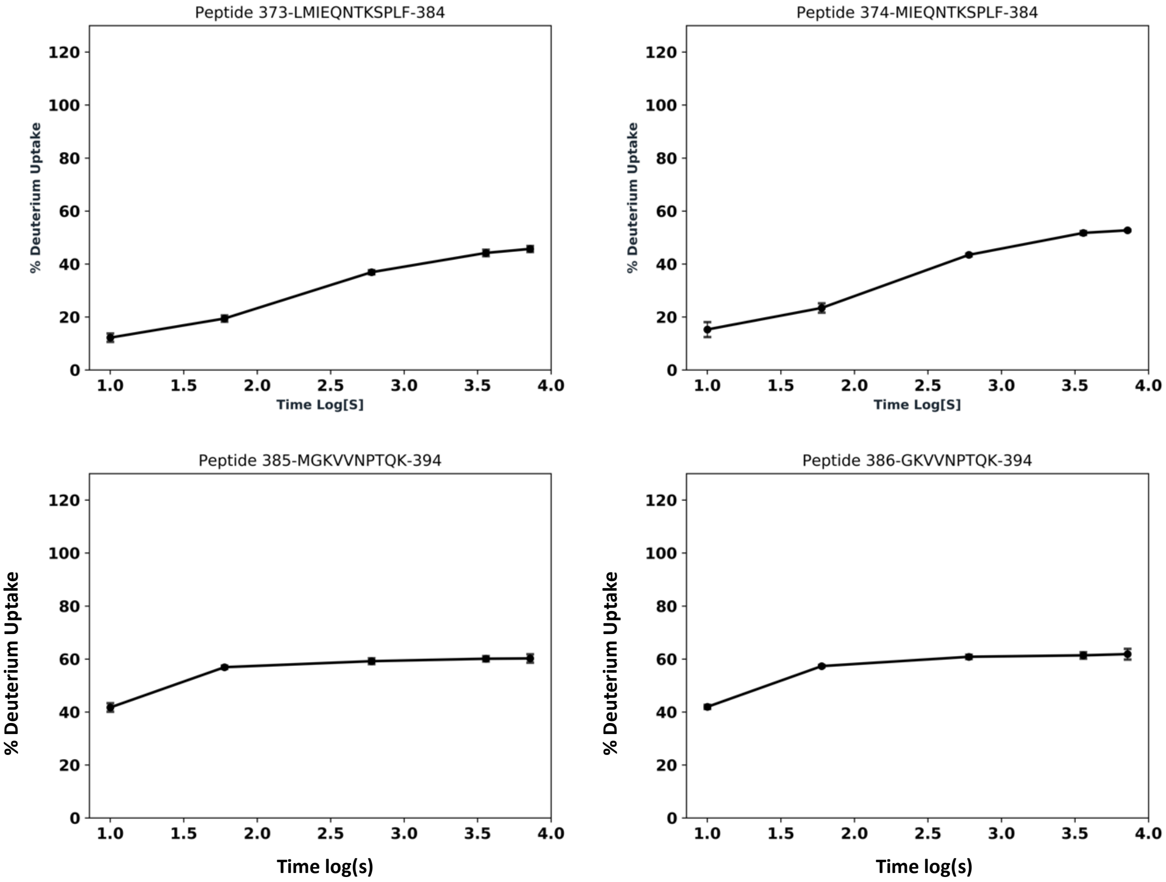


**Figure S7.** Kinetic uptake of deuterium for each peptide for each state with full length AAT in black, AAT190 biological replicate 1 in red and AAT190 biological replicate 2 in blue.

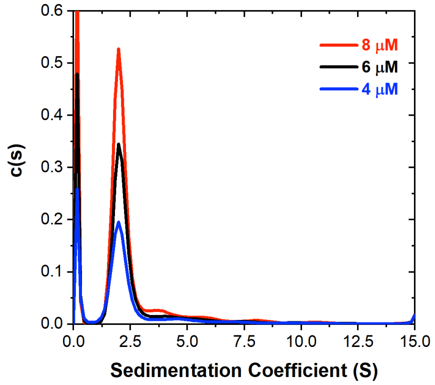


**Figure S8.** Distributions of sedimentation coefficients for 8 (red), 6 (black) and 4 (blue) µM of AAT190 prior to HDX-MS experiments indicating that this fragment was mainly monomeric. The AUC data were collected prior to the HDX-MS experiments the results of which are shown in Figure 4 in the main text. For all three distributions, the peak near 0 arises from a population of small molecules much smaller than the protein fragments.

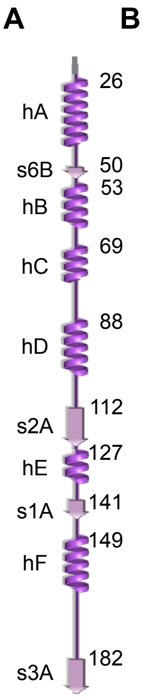


B

AAT190

| Peptide | 10 sec | 1 min | 10 min | 60 min | 120 min |
|---|---|---|---|---|---|
| 3-23 | 95.9 | 96.4 | 96.1 | 99.1 | 100.6 |
| 24-32 | 81.8 | 92.2 | 98.3 | 97.8 | 99.5 |
| 34-37 | 62.8 | 99.8 | 107.8 | 109.4 | 110.9 |
| 38-50 | 82.9 | 98.8 | 104.4 | 107.6 | 106.8 |
| 38-51 | 78.1 | 96.5 | 104.4 | 106 | 107.4 |
| 38-52 | 75.1 | 93.5 | 102.1 | 104 | 104.9 |
| 52-60 | 66.3 | 89.7 | 97.9 | 98.9 | 98.4 |
| 57-61 | 49.3 | 72.3 | 86.9 | 93 | 95.4 |
| 62-74 | 68 | 74.8 | 82.4 | 97.1 | 103.5 |
| 65-77 | 97.7 | 98.3 | 102.1 | 109.7 | 111.2 |
| 78-82 | 99.4 | 92.8 | 99.7 | 93.9 | 95.7 |
| 82-96 | 90.1 | 96.1 | 101.9 | 102.4 | 105.7 |
| 82-99 | 81.2 | 93.8 | 102.2 | 104.8 | 105.5 |
| 83-96 | 89.2 | 95.9 | 100.7 | 103.6 | 105.1 |
| 83-99 | 79.6 | 95.1 | 103 | 105.9 | 106.5 |
| 85-96 | 83.2 | 93 | 99.4 | 99.5 | 101.3 |
| 85-99 | 79.5 | 96.9 | 106 | 107.8 | 109.1 |
| 100-110 | 92.3 | 101.9 | 104.8 | 105.3 | 106.8 |
| 100-112 | 94.8 | 101.1 | 104.3 | 104.5 | 106.5 |
| 119-126 | 72.5 | 90.4 | 95.2 | 95.7 | 95.5 |
| 127-133 | 26.9 | 78.6 | 102.8 | 107 | 102.9 |
| 127-142 | 37.2 | 81.4 | 100.2 | 99.9 | 100.4 |
| 131-142 | 43.5 | 86 | 104.1 | 107 | 108.5 |
| 143-159 | 74.6 | 80.3 | 95.2 | 108.7 | 116.1 |
| 160-171 | 62.4 | 66.7 | 84.9 | 99.2 | 105.8 |
| 160-172 | 64.5 | 70.5 | 88.2 | 103 | 109.7 |
| 172-182 | 66.1 | 78.1 | 86.6 | 89.3 | 91.8 |
| 183-187 | 18.7 | 44.6 | 61.2 | 67.2 | 69.9 |

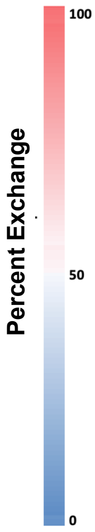


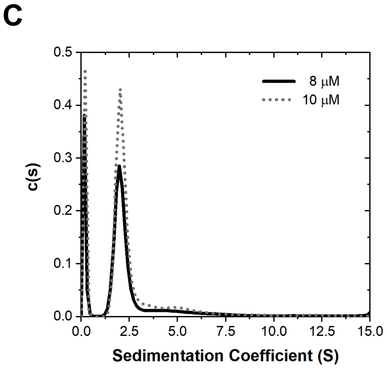


**Figure S9**. Protection from exchange over time for the second AAT190 biological replicate. (A) Schematic of the full-length AAT secondary structure. (B) Percent deuterium uptake for individual peptides at each incubation time point for AAT190. Peptide sequence numbers are listed in the column on the left and percent deuteration over time is colored from blue (0%) to red (100%). (C) Distributions of sedimentation coefficients for 10 (dotted) and 8 (solid) µM of AAT190 prior to HDX-MS experiments indicating that this fragment was mainly monomeric. For both distributions, the peak near 0 arises from a population of small molecules much smaller than the protein fragments.

A

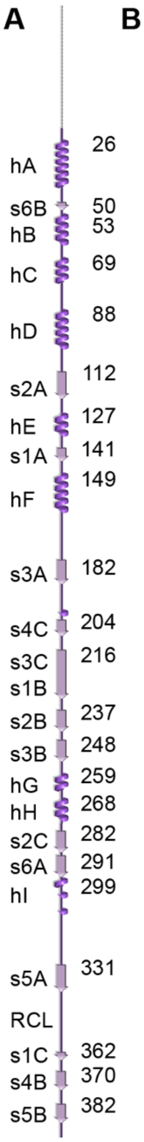


B

**Full-Length AAT**

| Peptide | 10 sec | 1 min | 10 min | 60 min | 120 min |
|---|---|---|---|---|---|
| 3-23 | 116.2 | 116.0 | 106.0 | 103.2 | 93.8 |
| 24-32 | 39.8 | 80.3 | 109.4 | 113.0 | 112.2 |
| 34-37 | 3.9 | 4.5 | 6.4 | 11.6 | 16.4 |
| 38-50 | 46.9 | 49.9 | 54.4 | 67.0 | 71.6 |
| 38-51 | 43.4 | 53.8 | 59.6 | 71.4 | 75.0 |
| 38-52 | 36.7 | 46.2 | 51.6 | 61.7 | 64.8 |
| 52-60 | 1.7 | 1.6 | 2.1 | 6.2 | 10.0 |
| 57-61 | 2.6 | 3.5 | 4.1 | 4.4 | 3.7 |
| 62-74 | 38.3 | 39.3 | 46.7 | 61.1 | 65.9 |
| 65-77 | 37.0 | 36.4 | 46.0 | 59.6 | 63.8 |
| 78-82 | 10.9 | 10.4 | 14.2 | 36.1 | 47.5 |
| 82-96 | 50.7 | 63.9 | 87.1 | 104.2 | 108.6 |
| 82-99 | 39.0 | 50.9 | 73.8 | 93.0 | 99.1 |
| 83-96 | 46.5 | 58.9 | 82.5 | 100.8 | 106.3 |
| 83-96 | 43.8 | 53.7 | 74.4 | 90.2 | 97.8 |
| 85-96 | 33.6 | 44.4 | 70.6 | 93.1 | 99.4 |
| 85-99 | 24.3 | 33.4 | 63.0 | 85.8 | 95.2 |
| 100-110 | 58.9 | 80.7 | 100.9 | 110.4 | 109.2 |
| 100-112 | 60.1 | 87.5 | 106.9 | 116.5 | 116.1 |
| 110-119 | 37.8 | 54.3 | 69.5 | 78.5 | 82.0 |
| 119-126 | 61.2 | 68.5 | 84.6 | 95.5 | 96.2 |
| 127-133 | 14.2 | 16.4 | 29.8 | 65.9 | 82.5 |
| 127-142 | 13.1 | 28.0 | 51.7 | 72.8 | 83.0 |
| 131-142 | 17.9 | 41.1 | 72.7 | 85.8 | 91.7 |
| 143-159 | 39.2 | 50.0 | 64.2 | 95.6 | 109.4 |
| 160-171 | 50.0 | 98.5 | 112.2 | 118.3 | 119.5 |
| 160-172 | 44.5 | 91.8 | 108.7 | 114.6 | 116.7 |
| 172-182 | 60.8 | 74.9 | 87.0 | 90.5 | 89.6 |
| 183-187 | 0.9 | 0.2 | 0.1 | 0.4 | 0.1 |
| 188-199 | 26.8 | 46.2 | 80.2 | 97.5 | 99.9 |
| 241-248 | 41.8 | 51.0 | 70.0 | 92.9 | 101.5 |
| 241-251 | 22.1 | 27.1 | 37.4 | 52.6 | 59.1 |
| 242-251 | 17.7 | 22.5 | 33.7 | 48.7 | 53.7 |
| 252-265 | 42.0 | 47.3 | 60.7 | 69.6 | 71.1 |
| 252-266 | 39.2 | 44.3 | 55.9 | 64.6 | 66.5 |
| 252-270 | 32.5 | 35.8 | 45.3 | 56.0 | 60.2 |
| 253-265 | 56.5 | 61.2 | 76.8 | 88.0 | 91.6 |
| 253-270 | 36.3 | 39.9 | 49.9 | 61.3 | 66.2 |
| 271-275 | 9.0 | 37.6 | 72.9 | 91.2 | 98.4 |
| 292-303 | 47.0 | 69.7 | 75.9 | 84.4 | 87.5 |
| 300-317 | 32.8 | 51.8 | 71.8 | 89.9 | 94.8 |
| 304-311 | 19.0 | 36.9 | 59.3 | 89.0 | 94.9 |
| 304-318 | 28.7 | 47.0 | 73.3 | 96.0 | 100.6 |
| 304-338 | 27.4 | 48.7 | 75.7 | 87.5 | 92.7 |
| 318-338 | 32.9 | 51.9 | 72.2 | 78.5 | 81.4 |
| 319-338 | 33.3 | 51.8 | 71.3 | 77.6 | 80.0 |
| 321-338 | 31.6 | 47.4 | 61.5 | 68.1 | 70.6 |
| 325-338 | 23.3 | 34.1 | 49.4 | 55.7 | 59.1 |
| 330-338 | 22.0 | 26.1 | 26.1 | 27.1 | 27.4 |
| 353-366 | 70.3 | 83.4 | 84.8 | 81.5 | 79.6 |
| 367-373 | 1.7 | 2.1 | 1.8 | 0.5 | 2.0 |
| 373-384 | 12.2 | 19.4 | 36.9 | 44.2 | 45.7 |
| 385-394 | 41.7 | 56.9 | 59.2 | 60.1 | 60.2 |
| 386-394 | 41.9 | 57.3 | 60.8 | 61.4 | 61.8 |

C

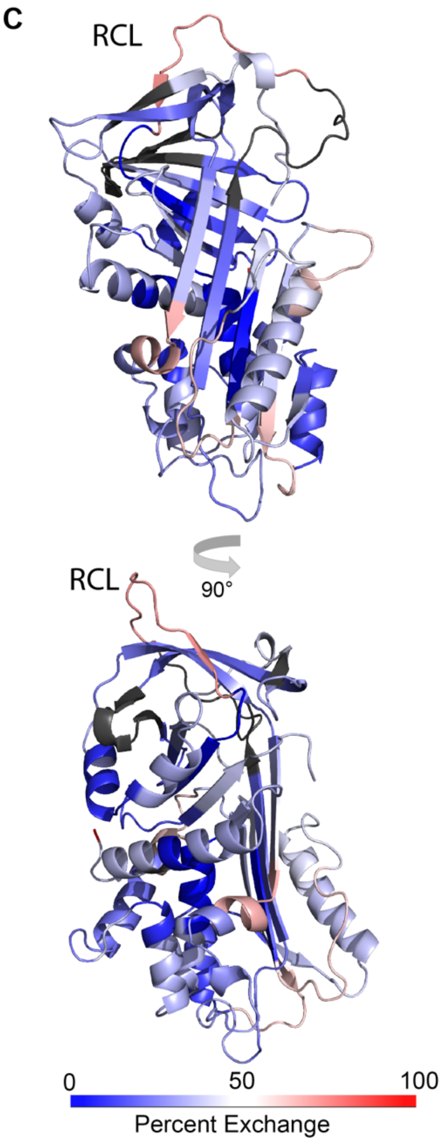


**Figure S10.** Protection from exchange for full-length AAT. (A) Schematic of the full-length AAT secondary structure. (B) Percent deuterium uptake for individual peptides at each incubation time point for full-length AAT. Peptide sequence numbers are listed in the column on the left and percent deuteration over time is colored from blue (0%) to red (100%). (C) Percent deuteration after 10 sec of incubation in $^2H_2O$ for full-length AAT (PDB: 1QLP (7) with the disordered N-terminus added using the CHARMM GUI (8)). Deuteration over time is colored from blue (0%) to red (100%).

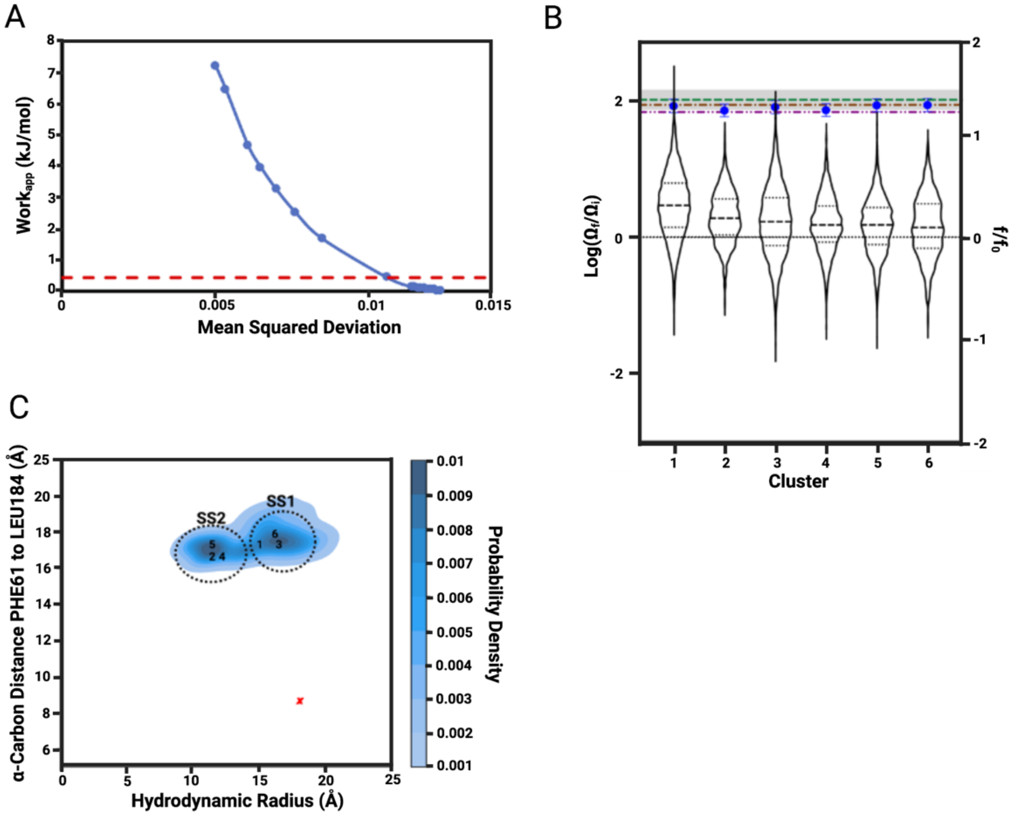


**Figure S11.** For AAT190 the HDXer method generates two sets of conformational ensembles and roughly predicts the frictional coefficient. (A) The decision plot obtained when reweighting the simulated tempering ensemble to the experimental AAT190 HDX-MS data using the HDXer approach (12, 13). The point of inflection in the decision plot of apparent work, $W_{app,}$ versus mean square displacement (MSD) shows the onset of overfitting, characterized as a rapid increase in $W_{app}$ for very little reduction in MSD. For AAT190 this rapid increase in $W_{app}$ is observed at 0.439 kJ/mol yielding a γ value of 2 shown by the red dashed line. Where γ helps prevent overfitting of the experimental HDX-MS data. (B) Violin plot for the six clusters ranked in order of the change in the ratio of the optimized weight, $\Omega_f$, to the initial weight $\Omega_i$ calculated for each frame in the cluster (left y-axis). The tails represent the minimum and maximum weight in the cluster, the thick dashed lines are the mean weight, and the thin dashed lines bracket 50% of the population. The frictional ratio ($f/f_0$) (right y-axis) was calculated for each cluster using HullRad (14) where the blue dot is the mean and the error bars are the standard deviation for all of the structures in each cluster. The dark red line is the value of $f/f_0$ calculated for the entire simulated tempering trajectory prior to reweighting and the gray line is the value of $f/f_0$ calculated from the 1 µs conventional MD simulation for AAT190 starting from the conformation in the x-ray crystal structure (PDB: 1QLP (7)). The experimentally determined value of $f/f_0$ is indicated by the green line with the 10% uncertainty indicated by the gray shading. (C) Kernel density estimation plot for all six significantly upweighted clusters as hydrodynamic radius (Å) versus the α-carbon distance between conserved shutter residues (15) F61 in helix B and L184 in strand 3A. The average value for each cluster is plotted as the number corresponding to its ranking. The red x is the average value for residues 1 to 190 in a 1 µs conventional MD simulation of full-length AAT. The grey circles delineate the two observed substates: substate 1, SS1, containing clusters 1, 3 and 6 which have more expanded conformations and SS2 containing clusters 2, 4 and 5 for which the conformations are more compact.

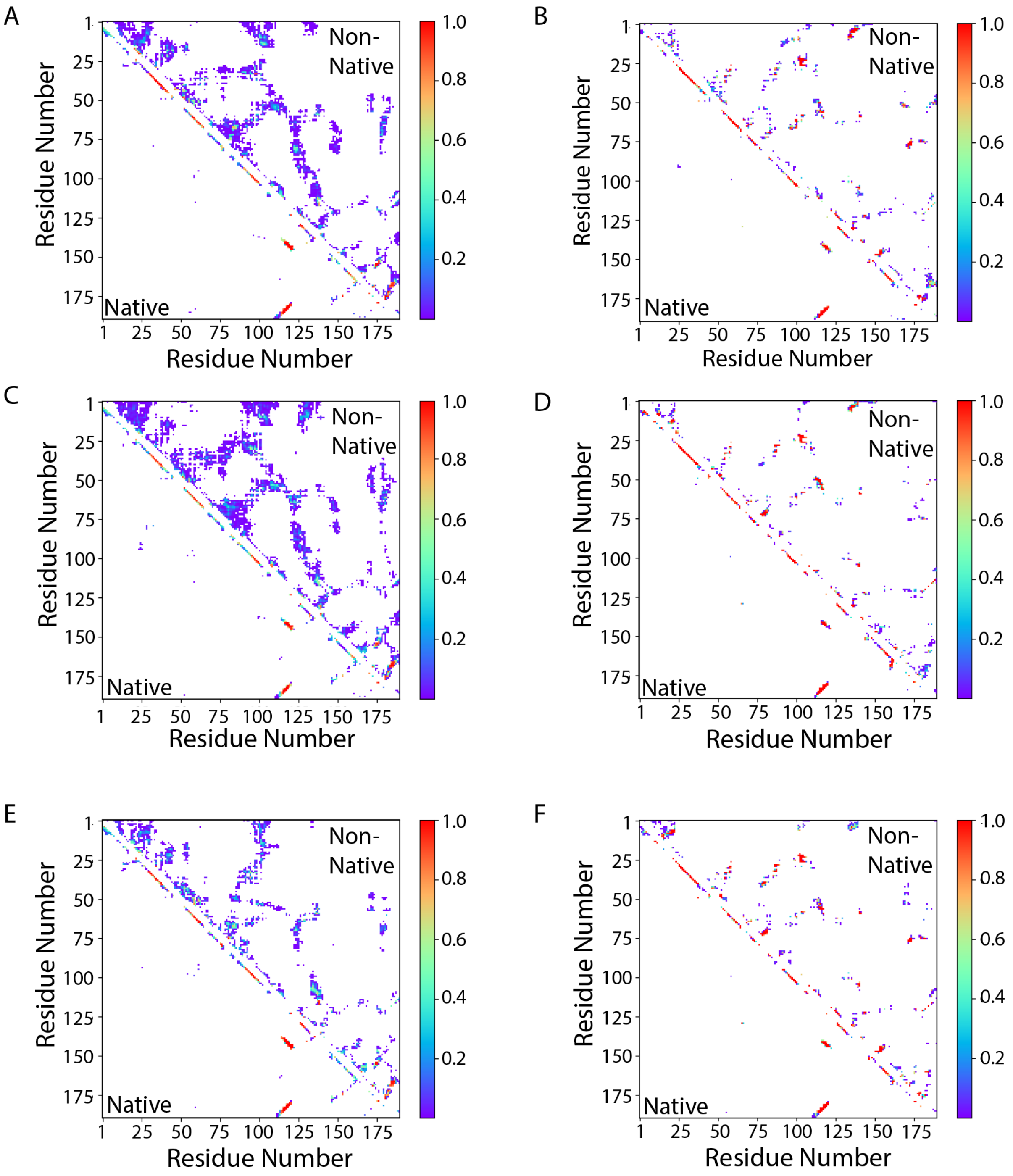


**Figure S12.** AAT190 contact maps for the six upweighted clusters from the HDXer analysis (12, 13) of the simulated tempering MD simulation. Clusters 1, 3 and 6 are shown in panels A, C and E, respectively. Clusters 2, 4 and 5 are shown in panels B, D and F, respectively. Native contacts that are present in the context of the full-length, active AAT structure are shown below the diagonal and non-native contacts are shown above the diagonal. The heat map indicates the relative occupancy determined using all of the frames in a given cluster (see Table S3 for the number of frames in each cluster). Contact maps were constructed using a cut-off of 7.5 Å for alpha carbons.

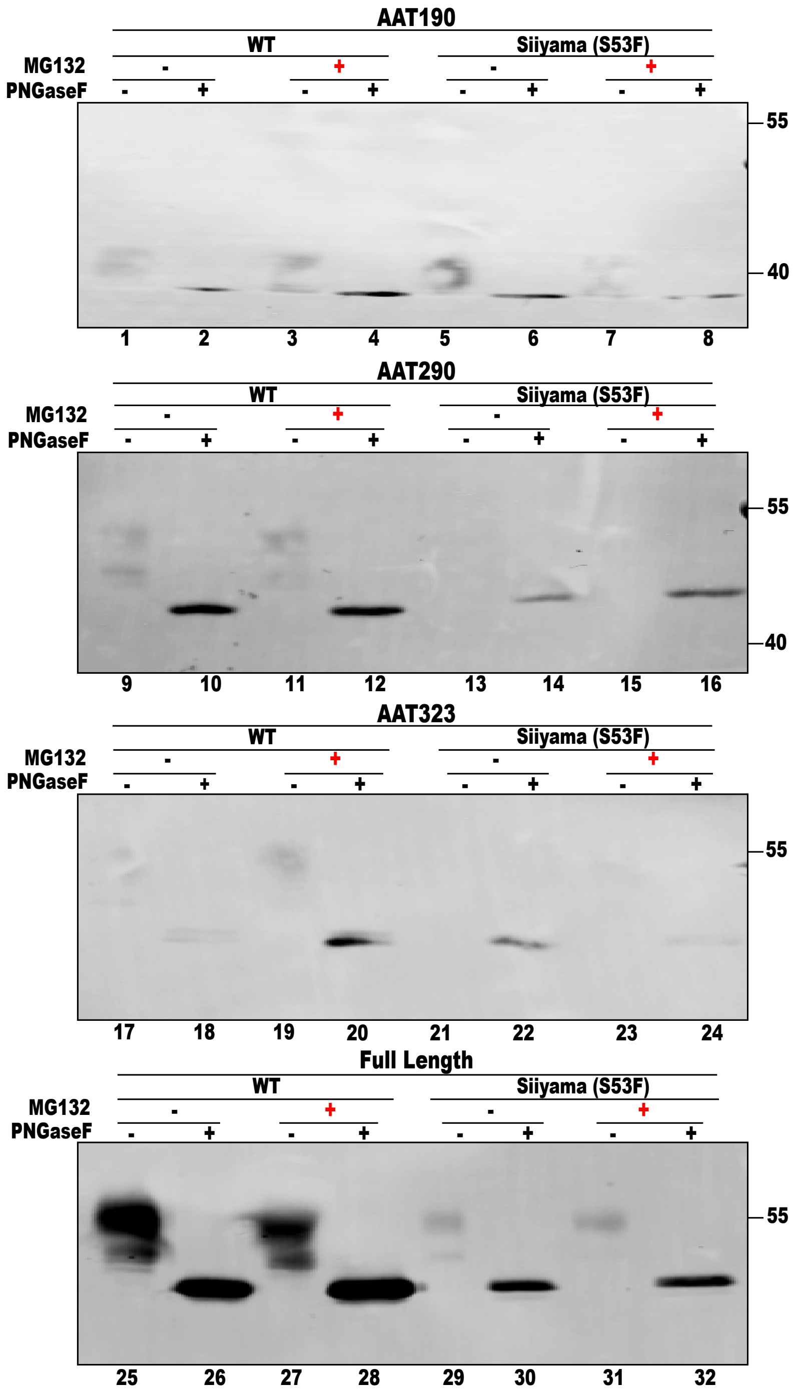


**Figure S13.** Full-length AAT and AAT fragments expressed in CHO cells are glycosylated. The cells were grown for 20 hr after transfection; then grown for a further 20 hr in the absence (black minus symbol) or presence (red plus symbol) of 20 µM of the proteasome inhibitor MG132. The cell media was collected, split in half and one half was treated with 500 units of the glycosidase PNGase F for 1 hr at 37° C (lower plus symbol). AAT constructs contained an N-terminal Myc tag (Table S2) and were detected using a Myc antibody. All constructs were glycosylated as evident from the shift to lower molecular weight upon treatment with PNGase F.

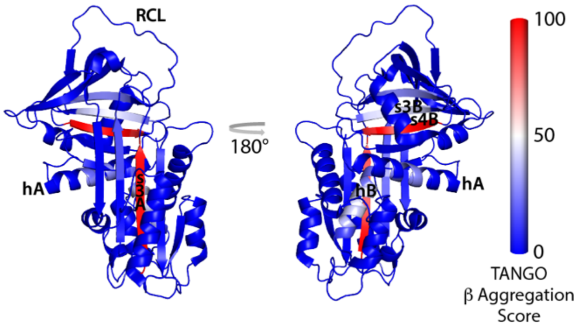


**Figure S14.** Strands 3A (residues 182-190) and 4B (residues 370-375) are predicted to be particularly aggregation prone. TANGO β aggregation scores (16, 17) mapped on the full-length AAT structure (PDB: 1QLP (7)) from low predicted β aggregation propensity (blue) to high (red).

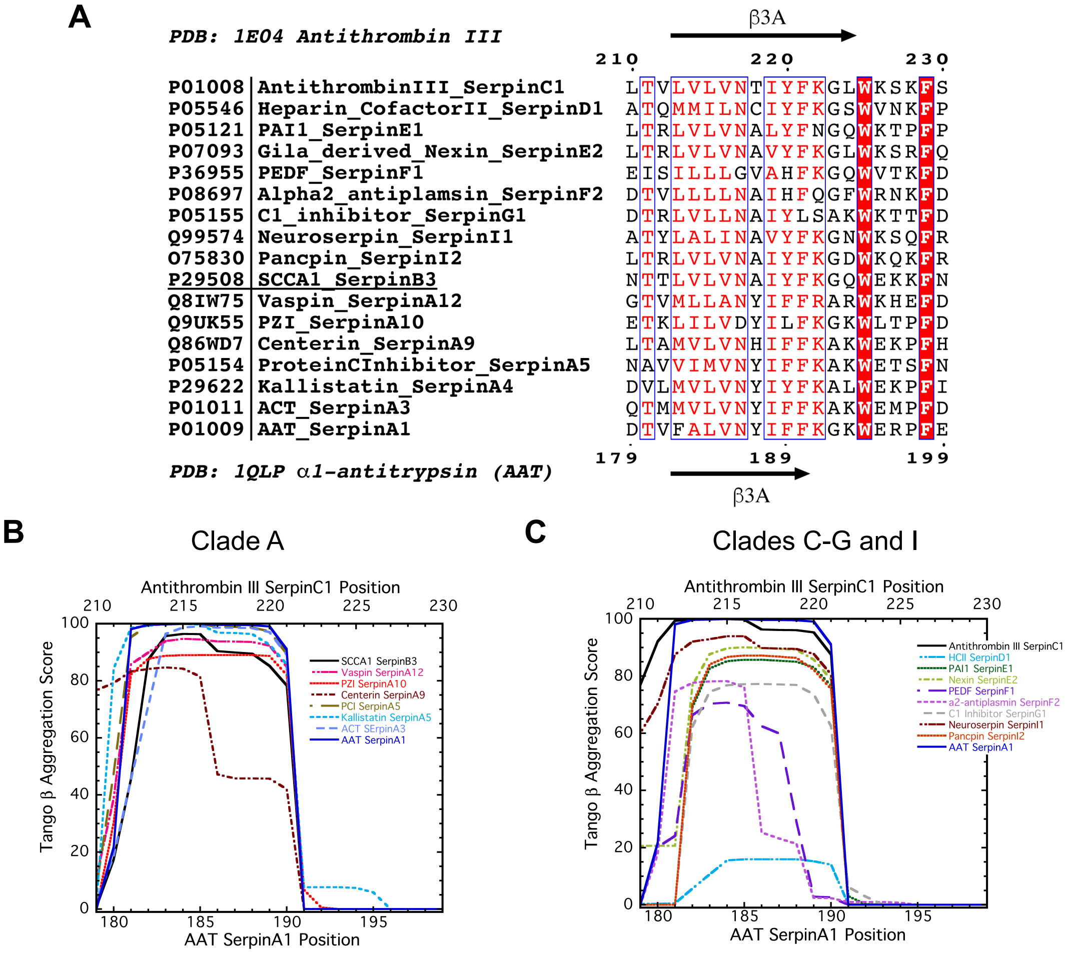


**Figure S15.** In human inhibitory serpins strand 3A is predicted to be prone to β strand mediated aggregation. (A) Clustal Omega (18) sequence alignment of the s3A region for human extracellular inhibitory serpins and a sample intracellular inhibitory serpin, squamous cell carcinoma antigen 1 (SCCA1, SerpinB3) with secondary structure assignments from antithrombin III (SerpinC1, PDB: 1E04 chain I (19)) and AAT (SerpinA1, PDB: 1QLP (7)) on the top and bottom of the alignment respectively. The Uniprot (20) identifier is listed for each serpin in the alignment. (B) Tango cross β-aggregation scores (16, 17) for human extracellular, inhibitory serpins from clade A. The intracellular inhibitory serpin SCCA1 (SerpinB3) is included for comparison. (C) Tango cross β-aggregation scores (16, 17) for human extracellular, inhibitory serpins from clades C-G and I. AAT (SerpinA1) is included for comparison. The colored alignment with the secondary structure was generated using ESPript 3.0 (21).

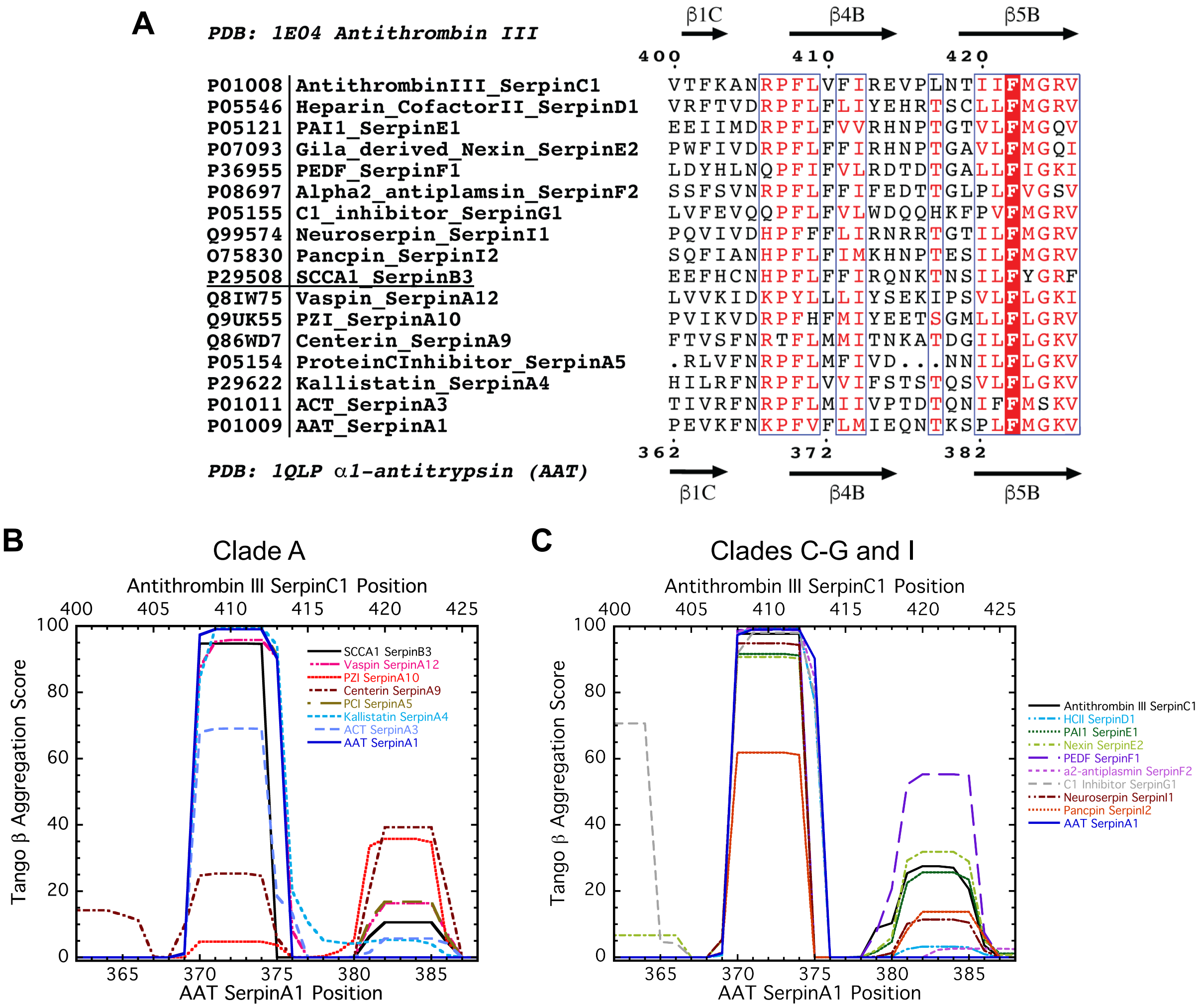


**Figure S16.** Strand 4B is predicted to be prone to β strand mediated aggregation in human extracellular inhibitory serpins. (A) The Clustal Omega (18) sequence alignment of the s4B/s5B region, which is near the serpin C-terminus, for human extracellular inhibitory serpins and a sample intracellular inhibitory serpin, squamous cell carcinoma antigen 1 (SCCA1, SerpinB3), is shown on top with secondary structure assignments from antithrombin III (SerpinC1, PDB: 1E04 chain I (19)) and AAT (SerpinA1, PDB: 1QLP (7)) on the top and bottom of the alignment respectively. The Uniprot (20) identifier is listed for each serpin. (B) Tango cross β-aggregation scores (16, 17) for human extracellular, inhibitory serpins from clade A. The intracellular inhibitory serpin SCCA1 (SerpinB3) is included for comparison. (C) Tango cross β-aggregation scores (16, 17) for human extracellular, inhibitory serpins from clades C-G and I. AAT (SerpinA1) is included for comparison. The colored alignment with the secondary structure assignment was generated using ESPript 3.0 (21). Note that in many of these serpins, strand 5B also has some predicted propensity for cross β-aggregation.

experiments with molecular simulations: Tutorials and applications of the HDXer ensemble reweighting software. *GitHub*. **Article v0.1**, https://github.com/TMB-CSB/hdxer_tutorials_livecoms
